# Multiple adaptive solutions to face climatic constraints: novel insights in the debate over the role of convergence in local adaptation

**DOI:** 10.1101/2021.11.18.469099

**Authors:** Badr Benjelloun, Kevin Leempoel, Frédéric Boyer, Sylvie Stucki, Ian Streeter, Pablo Orozco-terWengel, Florian J. Alberto, Bertrand Servin, Filippo Biscarini, Adriana Alberti, Stefan Engelen, Alessandra Stella, Licia Colli, Eric Coissac, Michael W. Bruford, Paolo Ajmone-Marsan, Riccardo Negrini, Laura Clarke, Paul Flicek, Abdelkader Chikhi, Stéphane Joost, Pierre Taberlet, François Pompanon

## Abstract

The extent to which genomic convergence shapes locally adapted phenotypes in different species remains a fundamental question in evolutionary biology. To bring new insights to this debate we set up a framework which aimed to compare the adaptive trajectories of two domesticated mammal species co-distributed in diversified landscapes. We sequenced the genomes of 160 sheep and 161 goats extensively managed along environmental gradients, including temperature, rainfall, seasonality and altitude, to identify genes and biological processes shaping local adaptation. Allele frequencies at adaptive loci were rarely found to vary gradually along environmental gradients, but rather displayed a discontinuous shift at the extremities of environmental clines. Of the more than 430 adaptive genes identified, only 6 were orthologous between sheep and goats and those responded differently to environmental pressures, suggesting different adaptive mechanisms in these two closely related species. Such diversity of adaptive pathways may result from a high number of biological functions involved in adaptation to multiple eco-climatic gradients, and provides more arguments for the role of contingency and stochasticity in adaptation rather than repeatability.

## Introduction

Local adaptation describes the adjustment or change in behaviour, physiology and morpho-anatomy of an organism to better fit its environment, and locally adapted populations are expected to exhibit higher fitness in their native habitats than populations from elsewhere (Kawecki and Ebert 2004). Identifying genomic changes underlying local adaptation is critical in addressing fundamental questions in evolutionary biology.

Most adaptive traits are affected by many segregating loci and are thus difficult to study (Ward and Kellis 2012), explaining why relatively few studies have so far elucidated fine scale patterns of local adaptation. For example, adaptation of deer mice to a sand coloured background is associated with independent selection on many Single Nucleotide Polymorphisms (SNP) within a single gene, each with a specific effect (Linnen et al. 2013). In contrast, a single mutation increases hair thickness, affects mammary fat pad size and increases eccrine gland number in mice (Kamberov et al. 2013). Such variation in the mechanisms and origins of adaptation suggests the occurrence of multiple evolutionary trajectories for any given trait, which may be divergent, parallel or convergent between populations or species (Elmer and Meyer 2011). Cases of adaptive convergence in phenotypic traits, biological functions, genes and even causal mutations are frequently reported in the literature, *e.g.* (Christin et al. 2010) (Losos 2011) (Elmer and Meyer 2011). Well-known examples are related to adaptive convergence to altitude in Tibetans and their domesticated animals, i.e. dogs, sheep, goats and falcons (Simonson et al. 2010) (Yi et al. 2010) (Zhan et al. 2013) (Gou et al. 2014) (Wei et al. 2016) (Song et al. 2016). However, other studies have reported similar adaptations based on divergent genomic architecture in closely related species/populations, such as a single pigmentation pattern controlled by different genes in beach mice (Steiner et al. 2009) (Manceau et al. 2010) or different genomic regions involved in the adaptation of co-existing stickleback species despite substantial phenotypic parallelism (Raeymaekers et al. 2017).

Since the Modern Evolutionary Synthesis, scientists have strived to distinguish the relative impact of stochasticity versus natural selection in speciation and diversification (Hey, J. 1998). This has led to the debate on whether evolution is unpredictable due to historical contingencies (Gould and Lewontin 1979) or repeatable (Conway Morris S 2003) or even whether both mechanisms are common. This has been invoked deeply in thought experiments such as “replaying life’s tape” (Gould S 1989). The debate over the role of convergence in local adaptation would help address these fundamental questions. Indeed, although adaptive convergence is overrepresented in the literature, this may be due to observation bias (i.e., more rarely published due to lack of interest). Thus, the extent to which genetic convergence is involved in shaping adaptive trajectories remains actually largely unknown (Riesch et al. 2018), necessitating whole genome studies on populations and species facing similar environmental pressures (Conte et al. 2012) to assess constraints acting on the evolution of genomes.

Previous studies addressing these key issues often compared isolated populations from contrasting environments under extremely divergent selective pressures (Gou et al. 2014) (Linnen et al. 2013). Moreover, environmental factors may vary gradually along gradients rather than changing abruptly, raising the question of whether genomic variants related to adaptation vary in a similar way (Riesch et al. 2018). A few studies have examined genomic signatures of selective pressures along environmental gradients including latitudinal clines in wing size of *Drosophila subobscura* (Huey et al. 2000) and flowering time in *Arabidopsis thaliana* (Stinchcombe et al. 2004). Landscape genomics (Manel et al. 2003) (Joost et al. 2007) is a well-suited approach to address such questions (Manel and Holderegger 2013). To date, most landscape genomics studies have used population genomics approaches to detect the adaptive variation within genomes in a spatial context (Manel and Holderegger 2013), however specific methods have also been developed that directly correlate allele frequencies with environmental gradients (Joost et al. 2007) (Frichot et al. 2013) (Stucki et al. 2017).

Here, we used a large-scale landscape genomics design, relying on an individual-based sampling and Whole Genome Sequencing (WGS) to examine how adaptive genomic variation changes spatially along environmental gradients and thus scrutinise convergence for adaptive trajectories in different species experiencing the same environmental heterogeneity. As convergent adaptations are expected to occur more frequently in closely related species (Conte et al. 2012), we aimed to assess the extent to which genes or biological pathways are reused independently to develop similar adaptations, using the closely related taxa of sheep *(Ovis aries)* and goats *(Capra hircus)* in the same environment. The *Ovis/Capra* divergence occurred around 6 Mya (Hassanin et al. 2012), and both domestic species experienced similar histories with overlapping centres of domestication around 10 kya ago near the Fertile-Crescent (Vigne et al. 2005) (Zeder et al. 2006) (Naderi et al. 2008), and the same colonization routes led to their rapid spread over the old-world. The populations studied here have been managed extensively with limited anthropogenic selection, and have accumulated valuable adaptations to their local environment (climate, ecology and husbandry), while maintaining a high level of genetic diversity (Taberlet et al. 2008). We focused on indigenous codistributed sheep and goat landraces in Morocco, and sequenced the genomes of 161 goats and 160 sheep representative of populations inhabiting Morocco’s high heterogeneity of geo-climatic conditions (e.g., Mediterranean, oceanic, high mountains, desert; Supplementary Fig. 1). We detected genomic signatures of selection associated with a wide variety of eco-climatic pressures that allowed us to identify sets of genes and biological processes that have shaped adaptation in sheep and goats. We characterized the patterns of variation in adaptive allele frequencies that exhibited variation along environmental clines. Finally, we showed that the genetic architecture underlying the adaptive responses to the same environmental gradients were mostly divergent between the two species.

## Results

### Genetic diversity and population structure

Whole genome sequences at 12X coverage were generated for 160 sheep and 161 goats and enabled the identification of more than 38.2 and 31.6 million variants, respectively. Among these variants, approximately 6.7% comprised small insertions/deletions (indels), and a low proportion (2.1% in sheep and 0.7% goats) of SNPs showed more than two alleles (mainly tri-allelic). Rare variants characterized by a minor allele frequency (MAF) of less than 5% comprised approximately 17 and 18 million SNPs in sheep and goats, respectively. Whole genome nucleotide diversity (*π*) was higher but not significantly so (*t-test*; p>0.05) in sheep than in goats. The average heterozygosity (*Ho*) was not significantly different between sheep and goats although the latter displayed a significantly higher inbreeding coefficient (*F*) (p<0.05; Table 1; see Methods).

**Table 1.**
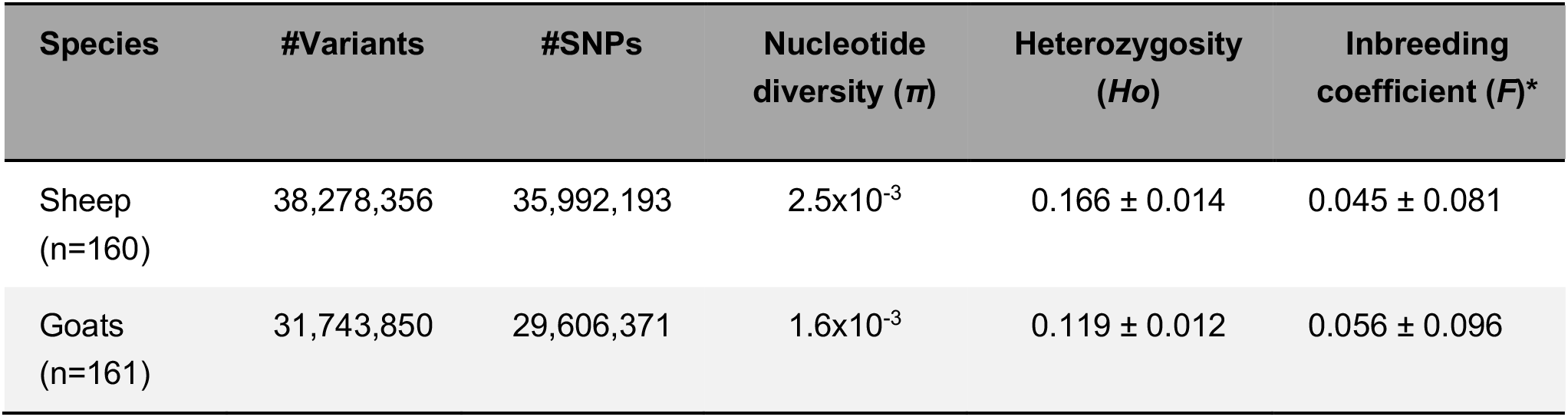
Whole Genome Diversity in Moroccan sheep and goats.

Genetic structure was analysed with sNMF (Frichot et al. 2013) using only biallelic SNPs (see Methods) and showed weak geographic patterns arising when setting three genetic clusters, with one cluster prevalent in the North for goats, and one cluster prevalent in the West for sheep (Fig. 1). However, cross-entropy analysis showed the most likely clustering corresponded to a single unit, suggesting no significant population structure (Supplementary Fig. 2 and Supplementary Fig. 3). This was consistent with PCA analysis where the first and second principal components showed no obvious population structure in either species, regardless of breed identification (Supplementary Fig. 4).

**Fig. 1.**
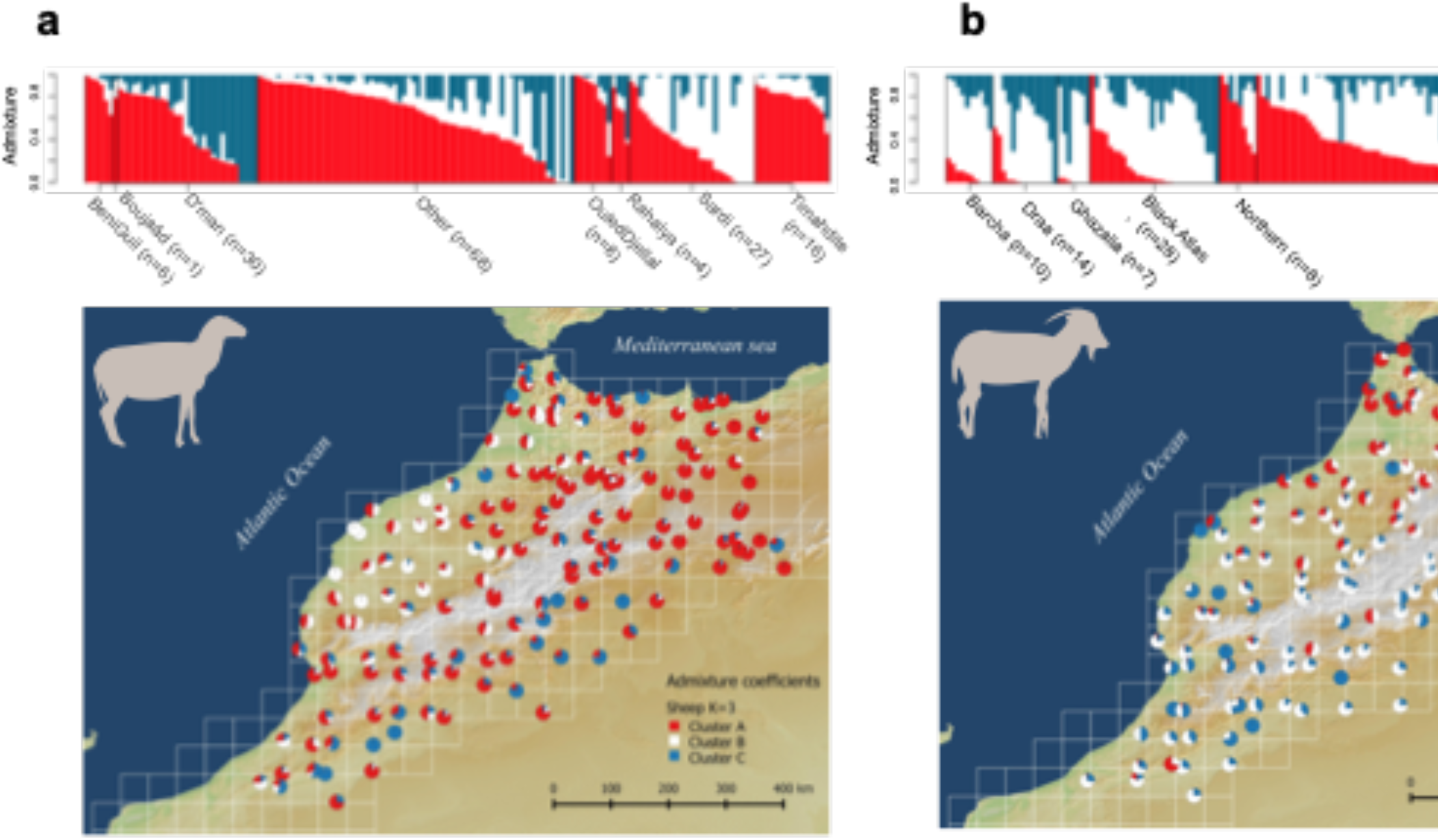
Genetic structure and sampling distribution for sheep (a) and goats (b). The upper charts show the proportions of genomes assigned to 3 genetic clusters according to the breed (i.e., admixture coefficients inferred by sNMF for the most likely number of clusters for both species). On the maps, each individual is represented by a pie chart located at the sampling site and showing the admixture coefficients.

### Signatures of selection

We investigated signatures of selection associated with the 10 least correlated environmental variables (with |Pearson’s r|<0.8) representing altitude, topography, solar radiation, temperature, rainfall, temperature seasonality and rainfall seasonality (see Methods and Table 2), using two approaches: 1) A correlative approach, Samβada (q-value threshold of 0.1; (Stucki et al. 2017), see Methods), was used to identify changes in allele presence/absence of biallelic SNPs along environmental variables using logistic regression for all individuals. In addition, a population-based approach was conducted by contrasting 2 groups comprising 20 individuals each from locations with the most extreme environmental values for each environmental gradient. We then calculated haplotype-based statistics (XP-CLR (Chen et al. 2010)) and single-locus *Fst* values (Weir and Cockerham 1984) along the genome, and defined SNPs among the top 0.1% XP-CLR scores and the top 0.5% *Fst* values as outlier variants (see Methods).

**Table 2.** Number of outlier variants and related genes identified for each environmental parameter in sheep and goats. The number of outliers detected via the correlative approach (Samβada), the population based approach (XP-CLR/Fst) and both.

| Environmental parameter | WorldClim and DEM derived Variables | correlative approach | Number of variants/genes |  |  |  |  |
| --- | --- | --- | --- | --- | --- | --- | --- |
|  |  |  | Sheep population based approach | Common | correlative approach | Goat population based approach | Common |
| Altitude | **Altitude** (m) | 0/0 | 650/14 | - | 80/6 | 587/30 | 3/1 |
| Slope | **Slope** | 0/0 | 672/8 | - | 0/0 | 440/19 | - |
| Solar radiations | **ti2112**<br>(sunshine duration on December 21th) | 0/0 | 1066/25 | - | 0/0 | 452/20 | - |
| Temperature | **tmin8**<br>(average monthly minimum temperature in August) | 0/0 | 1262/39 | - | 0/0 | 585/16 | - |
|  | **tmean7**<br>(average monthly mean temperature in July) | 276/13 | 1653/27 | 173/7 | 0/0 | 427/17 | - |
| Rainfall | **Prec4**<br>(average monthly precipitation in April) | 839/32 | 1500/23 | 290/5 | 5/0 | 718/17 | 0/0 |
| Temperature seasonality | **bio7**<br>(temperature annual range) | 141/8 | 904/25 | 0/0 | 135/8 | 466/16 | 2/0 |
|  | **bio8**<br>(mean temperature of the wettest quarter) | 0/0 | 1145/21 | - | 0/0 | 421/19 | - |
|  | **bio3**<br>(isothermality) | 0/0 | 1203/33 | - | 0/0 | 567/26 | - |
| Rainfall seasonality | **bio15**<br>(precipitation seasonality, coefficient of variation) | 0/0 | 785/8 | - | 127/14 | 797/26 | 51/1 |
| Overall (association/variant genomic-region/gene) |  | 1242 / 758 | 10840/9434 | 463/317 | 347/311 | 5460/5292 | 56/56 |
|  |  | 108 / 41 | 210/183 | 4/8 | 61/16 | 297/198 | 5/2 |

Samβada identified 1,242 significant allele/environment associations involving 758 SNPs from 108 genomic regions (defined by adjacent outlier variants closer than 100kb) in sheep (Table 2, Supplementary Fig. 12). Of these SNPs, 93% clustered in 15 regions on 10 chromosomes, each region bearing at least 10 significant associations. SNPs belonging to the same cluster were associated with one or more environmental variables and displayed similar spatial patterns of allelic variation. Most of the genomic regions were associated with rainfall (prec4), temperature (tmean7) and annual temperature range (bio7). The largest region was centered on the *MC5R* gene on chromosome 23 and carried 839 significant allele/environment associations related to 383 SNPs that were mainly associated with rainfall (prec4), and to a lesser extent with temperature (tmax4, tmean7 and bio7). In goats, Samßada identified 497 significant allele/environment associations for 311 SNPs from 61 genomic regions (Table 2, Supplementary Fig. 12). Most associations were with annual temperature range (bio7), rainfall seasonality (bio15) and altitude. As with sheep, >90% of the significant SNPs were clustered, with 7 regions containing more than 10 significant associations. We observed a major cluster of 50 outliers mainly associated with rainfall seasonality (bio15) and altitude, centred on a non-annotated gene on chromosome 6. A search for its orthologue using Ensembl showed a high similarity (99.62% match) with the bovine gene *CAMK2D,* which is associated with the development of mammary tissue (Nguyen and Shively 2016) and cardiac function (Little et al. 2007). Four additional regions on chromosome 4, 24, 13 and 27 were associated with rainfall seasonality (bio15), and four regions on chromosome 4, 6, 9 and 11 were associated with temperature seasonality (bio7).

The population-based approach (XP-CLR/Fst) identified 9,433 outlier variants in sheep from 210 genomic regions, which were associated with 183 genes. Fourteen candidate genes were associated with altitude, 24 with rainfall (prec4), 25 with annual temperature range (bio7), 27 with temperature (tmean7) and 8 genes with slope (Table 2, Supplementary Fig. 13). In goats, we found 5,292 variants from 297 genomic regions linked to 198 genes. Thirty candidate genes were associated with altitude, 17 with rainfall (prec4), 16 with temperature annual range (bio7), 17 with temperature (tmean7) and 19 with slope (Table 2). Outlier loci associated with given environmental variables clearly showed an allele more prevalent under extreme environmental conditions as illustrated in Fig. 2C.

**Fig. 2.**
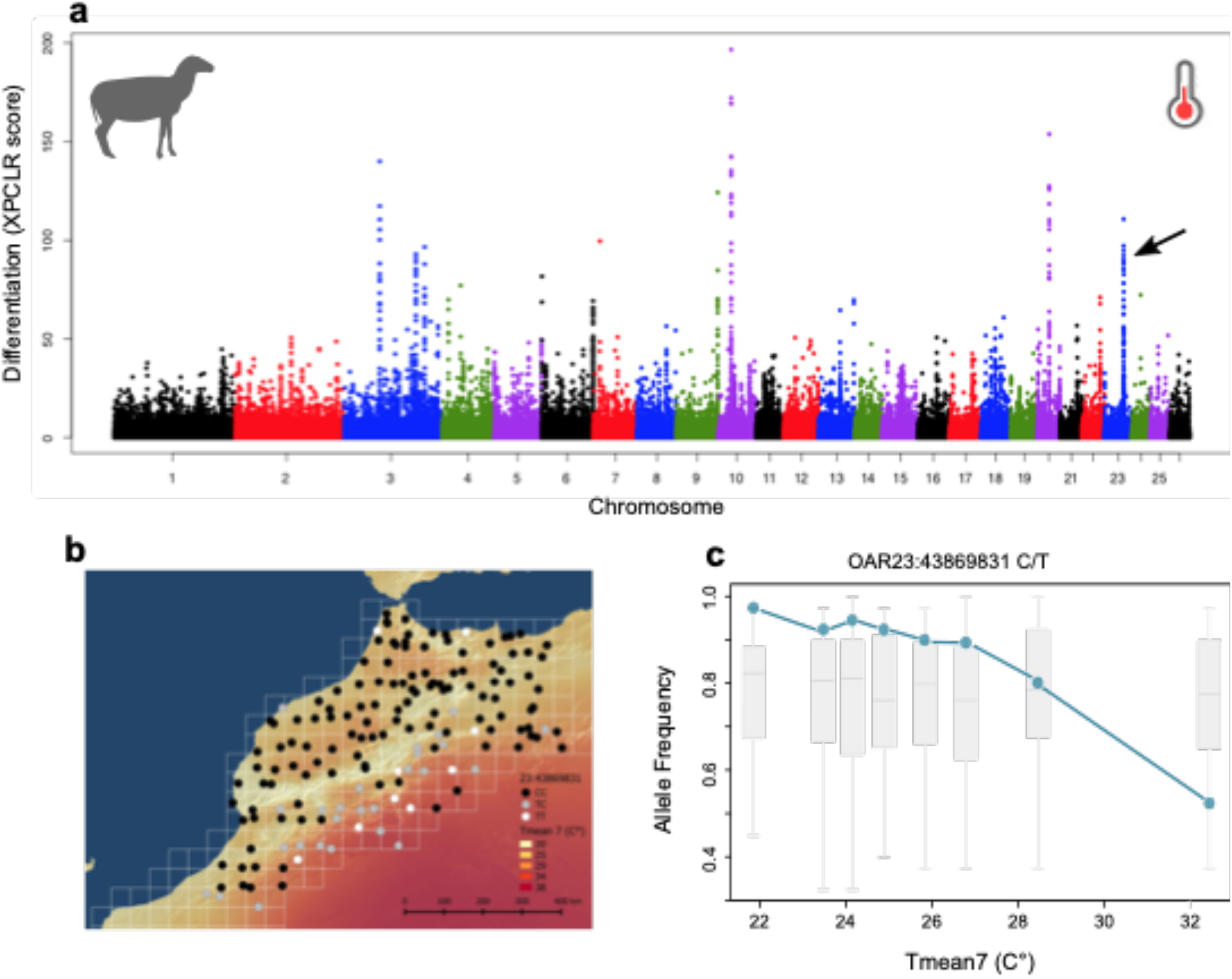
Genetic differentiation associated with temperature in sheep. (a) Manhattan plot of XP-CLR scores across autosomes in relation to the mean temperature of July (tmean7). The peak on chromosome 23 (arrow) comprises a missense SNP (OAR23:43869831 C/T). (b) Geographic variations of genotypes at this locus reported on the map of tmean7 variations. (c) Variation of allele frequency at this locus for groups of 20 sheep along the temperature gradient. The grey box-plots represent neutral variations of allelic frequencies for 100 variants randomly selected over the genome.

In total we detected a total of 9,875 and 5,547 outlier variants with at least one of the two approaches (324 and 56 common to both methods), corresponding to 291 and 346 genomic regions (4 and 5 regions common to both methods, Table 2) in sheep and goats, respectively. These regions are distributed over all chromosomes (see Supplementary Fig. 5-7). Among these regions, 19 were longer than 200 kb in sheep (largest 648 kb) with 16 such loci in goats (largest 427 kb). A total of 217 genes in sheep and 213 genes in goats were associated with at least one environmental variable (Supplementary Table 3). Among these genes, seven were detected by both methods in sheep, related to rainfall (prec4 and prec8) and temperature (tmean7 and tmax4). Among them, *MC5R* on chromosome 23, showed 2 missense alleles, which were present in the driest zones of the sampling area (Fig. 2C). In goats, two genes were detected by both methods, i.e., *CDH2,* located on chromosome 24, associated with rainfall seasonality (bio15) and *PXK* on chromosome 22, associated with altitude.

Most of the variants detected were intergenic (63% and 67% for sheep and goats, respectively) and intronic (27% and 25% for sheep and goats, respectively). Sixty-eight outlier SNPs were exonic (21 missenses and 47 synonymous) in sheep, with only one (a single missense mutation) in goats (Supplementary Table 3). These proportions were similar to that of the entire genome for sheep but there were more intronic and intergenic variants among outliers for goats (Chi-square test, P<0.05).

### Patterns of allelic variation at selected loci

We used an exploratory approach to evaluate genomic responses to environmental variation by characterizing the variation of allele frequencies along environmental variables for the most differentiated SNP locus within each selected genomic region for the 10 environmental parameters (Fig. 3A and Supplementary Fig. 8-11). We categorized each variation profile as gradual (i.e., linear) or punctual at one or both extremities of the gradient (Fig. 3A & 3B). For both species, all variation patterns were observed, with 69% of outliers in sheep and 80% of outliers in goats corresponding to punctual shifts occurring at extreme environmental values (i.e., low, high or both, *e.g.* Fig. 3 and Supplementary Fig. 8-11).

**Fig. 3.**
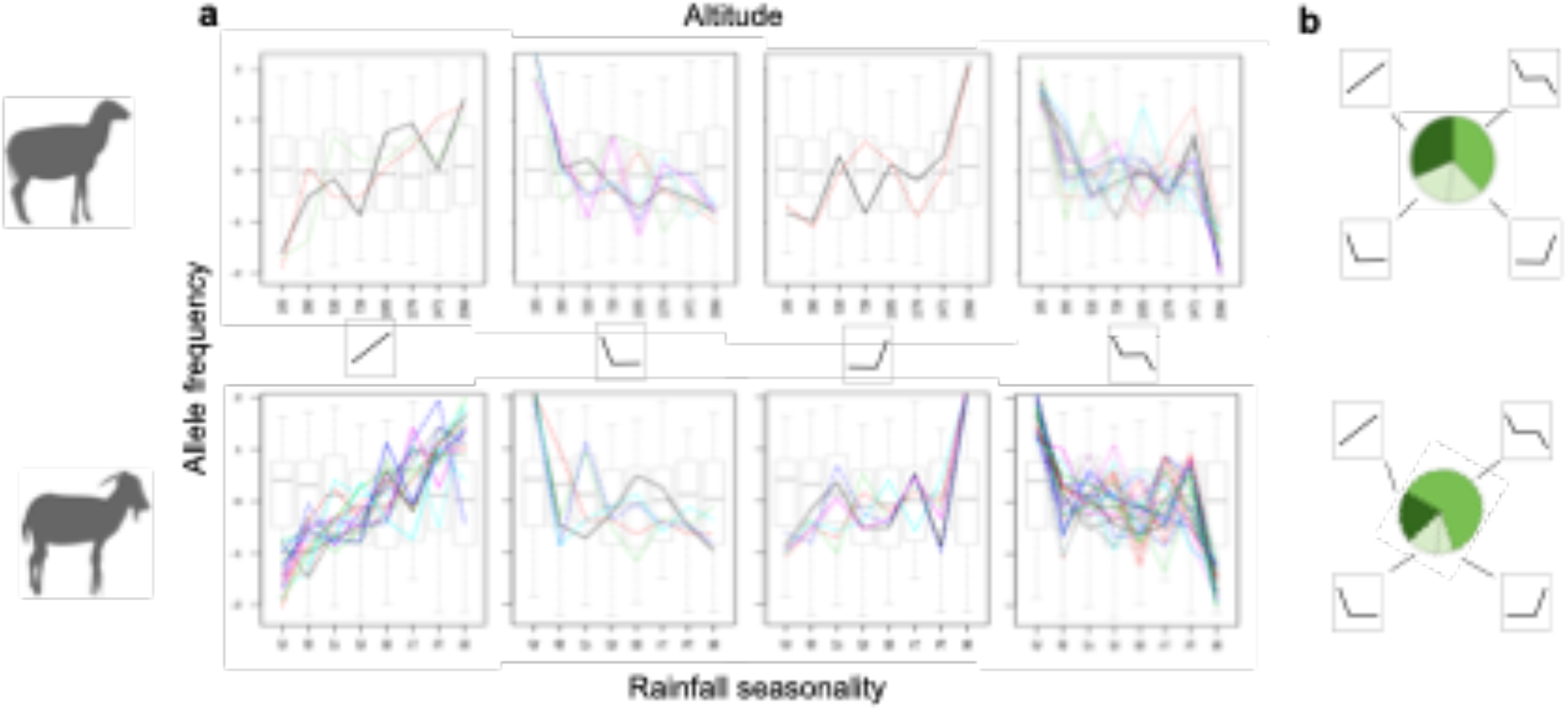
Variations of allele frequencies for candidate variants related with altitude in sheep (upper charts) and rainfall seasonality in goats (lower charts). (a) For each SNP, the variation profile of allele frequency (centred values) is assigned to one of the following categories: linear variation, uniform except at one extreme of the gradient, uniform except at both extremes (see methods). The grey box-plots represent neutral variations of allelic frequencies for a set of 100 random variants. (b) Proportions of variants assigned to each type of profile.

### Gene Ontology enrichment analysis

We performed Gene Ontology (GO) enrichment analyses with GOrilla (Eden et al. 2009) followed by a clustering with REVIGO (Supek et al. 2011) using the entire set of candidate genes (see Methods). In sheep, four GO categories were enriched in genes associated with altitude, including two GO terms linked to lung development related to the gene *NFIB*. Fourteen GO terms were enriched in genes associated with temperature (tmean7) and annual temperature range (bio7), depending on one main category related to the regulation of cAMP biosynthesis. One GO term was associated with temperature (tmin8), temperature seasonality (bio15) and solar radiation (ti2112) (Supplementary Table 5). For goats, 24 GO terms were enriched in genes associated with temperature seasonality (bio8), in five distinct categories, i.e. stem cell division, regulation of hydrolase activity, cellular component morphogenesis, locomotory behaviour and neuromuscular process. In addition, 21 GO terms associated with temperature (tmin8) corresponded mainly to the regulation of transmembrane cellular transport. We also found four GO terms associated with isothermality (bio3) and related to the cell response to superoxide and mitochondrial transport. Two GO terms were associated with rainfall seasonality (bio15). Slope, annual temperature range (bio7), rainfall (prec4) and solar radiation (ti2112) were associated with one GO term each (Supplementary Table 6).

### Orthologous genomic regions related to environmental variations in sheep and goats

We mapped all variants identified in sheep to the goat genome and *vice-versa* but found no orthologous outlier SNPs between species. Further, of the 424 putatively adaptive genes in sheep and goats, only six were orthologous (i.e., *BICC1, CA2, DENND4C, DUSP2, FAT3, TCP11L2;* Supplementary Table 3). Additionally, only a single genomic region was orthologous in both species where 26 variants associated with winter sunshine duration (ti2112) in sheep were within 100kb of 13 variants identified for the same variable in goats. Among the above genes, *CA2 and BICC1* were the only genes to be associated with the same variables (i.e., solar radiation ti2112 for *CA2* and rainfall in April prec4 for *BICC1*) in both species. *CA2* had one downstream outlier variant in goats and four intronic, three downstream and six upstream outlier variants in sheep. In contrast, *BICC1* had 24 intronic outlier variants in goats and 2 intronic variants in sheep.

## Discussion

### Performance of the experimental design for assessing convergent adaptive trajectories

Our study was specifically designed to assess the diversity and uniqueness of the adaptive response at two levels: (i) within-species by characterizing patterns of adaptive genetic variation along environmental gradients, (ii) between-species by comparing the adaptive responses of two closely related species along the same environmental gradient. Analysing the adaptive response of genomes along environmental gradients has mainly been studied via discrete sampling in contrasting extreme environments (Riesch et al. 2018), leaving large parts of these gradients unexplored. Our sampling strategy maximized the coverage of multiple environmental gradients by sequencing 160 whole genomes per species across an environmentally heterogeneous region of approximately 400,000 km^2^. As a consequence, the environmental clines explored could not be superimposed on latitudinal gradients or colonization routes, which are potential sources of confounding effects. We found high genomic variation in both species (38.6 and 31.7 million SNPs in sheep and goat, respectively), weakly structured over the area studied and among breeds (Fig. 1, Supplementary Fig. 2, Supplementary Fig. 3). This suggests that no strong bottlenecks have occurred during breed evolution and that gene-flow has been maintained (Benjelloun et al. 2015) and/or related to successive waves of colonisation along the Mediterranean routes of domestication (Pereira et al. 2009). This weak population structure has a clear advantage when looking for selective sweeps due to local adaptation, allowing us to avoid confounding demographic factors (Holderegger et al. 2008).

### Genetic differentiation along environmental gradients

Changes in adaptive allele frequencies in relation to the spatial variation of their relative fitness are usually expected to be either linear or sigmoidal (Novembre and Di Rienzo 2009), corresponding to patterns of either gradual or punctual genetic variation, respectively. The combination of methods used here (i.e. Samβada and XP-CLR) should in principle allow the detection of both types of patterns. We detected putatively adaptive genomic regions showing either a gradual or more frequently punctual frequency variation along multiple environmental gradients (69% in sheep and 80% in goats). This may be explained by stronger selection at environmental extremes, in accordance with the expectation that extreme values of environmental parameters are responsible for higher functional stress and thus stronger selective pressures (Grant et al. 2017). Even if we cannot exclude a lower power for the first approach as noted by Cao et al. (2021), this would also explain that Samβada identified substantially fewer SNPs than XP-CLR/Fst.

### Targets of selection and adaptive pathways

Provided that the spatial patterns of allelic frequencies reflect variation in adaptive phenotypes (Novembre and Di Rienzo 2009), identifying locally adapted loci and their associated environmental drivers is a first step towards understanding the genetic architecture underlying adaptive traits and the environmental conditions responsible for population divergence (Tiffin and Ross-Ibarra 2014). We identified genomic regions mainly responding to average temperature and rainfall, and to a lesser extent to seasonality, altitude, solar radiation, and topography. Most of the selective sweeps identified affected only non-coding variants as reported for other species, *e.g.* (Carneiro et al. 2014) (Ai et al. 2015) (Boitard et al. 2016). Functional annotation of non-coding variants is challenging as the gene(s) they affect can be located far away and might not even be the physically closest one. When located close to genes, they could be influencing transcription (Ward and Kellis 2012). Interestingly, Naval-Sanchez et al. (Naval-Sanchez et al. 2018) showed that domestication and subsequent selection have preferentially impacted proximal gene regulatory elements in sheep. Non-coding variants could also constitute a part of sequences regulating translation, stability and localization, *e.g.* (Chan et al. 2010) (GTEx Consortium 2017). Moreover, they could contribute more generally to functional regions that are not closely located to genes under regulation such as regulatory non-coding RNAs (e.g, miRNAs) (Noonan and McCallion 2010) (Dunham et al. 2012) (Johnson and Voight 2018). These observations are in accordance with theories suggesting that evolution may be facilitated more through changes in gene regulation than through changes in gene sequence (Carroll 2008). The current lack of knowledge on the identification and role of regulatory sequences in non-human species limits the interpretation of selective sweeps in adaptation to eco-climatic pressures and emphasises the need for a comprehensive and integrated resource on functional elements of animal genomes such as the FAANG initiative (https://www.animalgenome.org/community/FAANG/index).

We identified multiple gene-bearing regions affected by selective sweeps and searched for insight in their functional meaning by analysing enrichment of GO terms. In sheep, adaptation to altitude involves GO terms related to the growth and differentiation of lung cells whose proliferation is related to altitude (Heath et al. 1976) and hypoxia (Uhlik et al. 2005), and GO terms related to altitude also include heart rate regulation via the gene *TPM1.* More specifically, the gene *MCM3* was a top-candidate gene related to altitude. It regulates the activity of *HIF-1* (hypoxia-inducible factor 1) (Hubbi et al. 2011) (Semenza 2011), which has been shown with its paralog *HIF-2* (*EPAS1*) to be under selection at high altitude in several species including sheep and goats (Beall et al. 2010) (Simonson et al. 2010) (Yi et al. 2010) (Gou et al. 2014) (Wei et al. 2016) (Pan and Shen 2017). Even if the environmental pressures related to altitude encountered here (highest altitudes sampled at 2,300 m with likely transhumance up to 3,300 m ASL) might differ from that of previous studies targeting higher altitudes (e.g. Tibetan populations, > 3,000 m ASL), our findings illustrate that different genes of the same pathway might be involved in different adaptive responses to similar environmental pressures (Fig. 4B). In goats, the GO term “Regulation of respiratory gaseous exchange” (GO:0043576) was related to temperature annual range via the genes *PASK* and *GLS.* This is consistent with a thermoregulation via panting in goats experiencing highly fluctuating temperatures (Dmiel and Robertshaw 1983); (Baker 1989). Indeed, goats have two different thermoregulation mechanisms, panting or sweating, according to the level of dehydration (Dmiel and Robertshaw 1983) (Baker 1989) depending on the breed (Dmiel et al. 1979). This result might be partly driven by the presence of Draa goats in our samples (7 Draa among the 20 goats in the group with the highest annual temperature range) even in the absence of breed-specific genetic clustering. The same GO category has already been shown to be enriched for selected genes in this breed which is adapted to oases and desert areas in Morocco (Benjelloun et al. 2015).

**Fig. 4.**
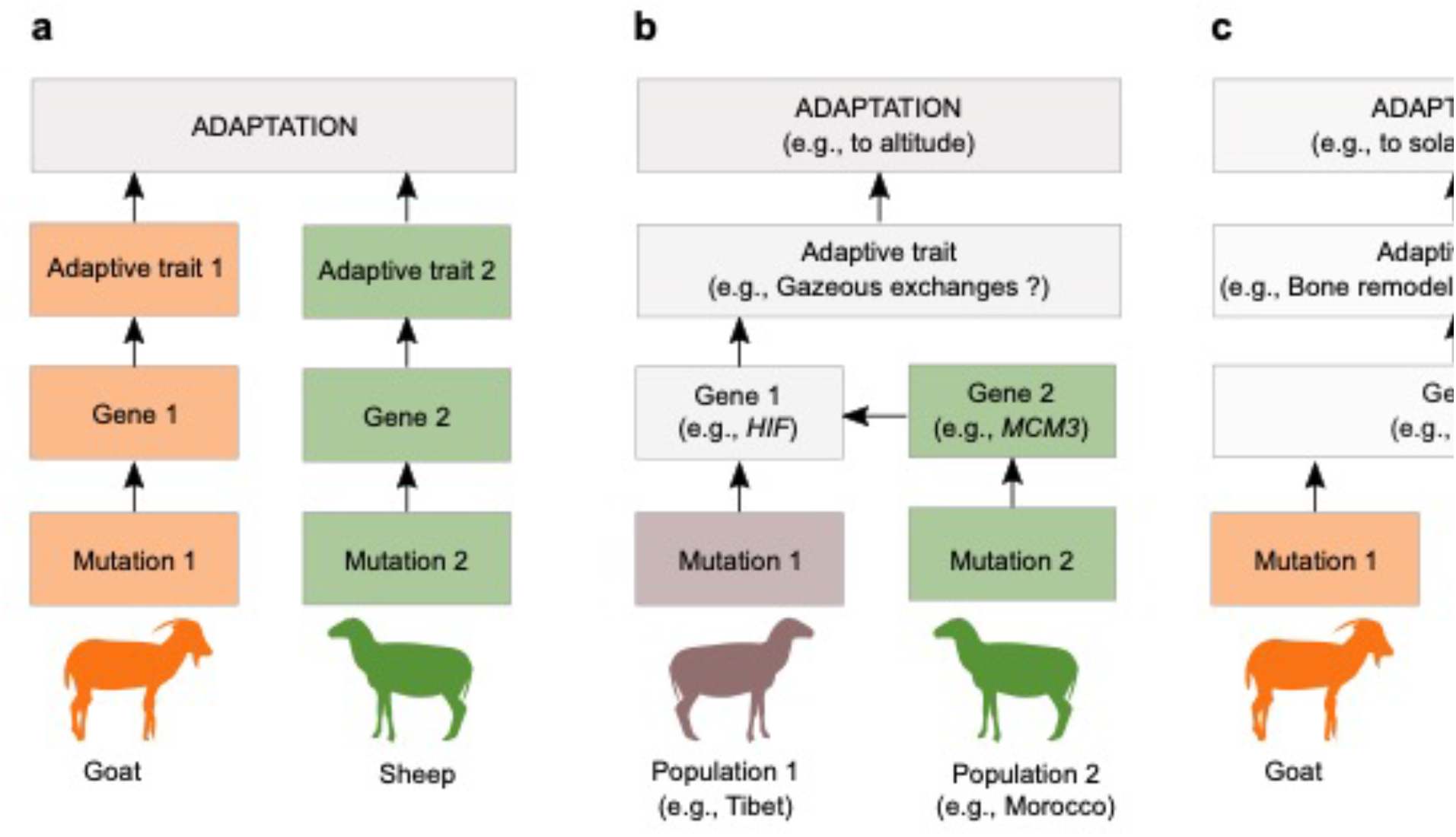
Different levels of similarity between independent adaptations to the same environmental pressure. (a) Variations at different locus involving different biological processes is the general case in this study. (b) Variations at different locus involved in the same biological process through e.g., regulation interactions. This is illustrated in sheep by *MCM3* related to altitude in this study which regulates *HIF* paralogs under selection in other sheep populations (Wei et al. 2016). (c) Variations at the same locus through different mutations (e.g., *CA2* associated with solar radiation). The adaptive traits are hypothesized based on the GO terms enriched in candidate genes.

GO terms linked to cellular component morphogenesis and stem cell division and differentiation were associated with temperature seasonality. Among the outlier genes identified, *PAFAH1B1* is involved in stem cell morphogenesis, *ABL1* in B cell differentiation, *NOTCH1* in angiogenesis (Scott et al. 2015) and *ALS2* in several cellular processes, e.g. endocytosis, development of axons and dendrites (Ratti and Berry 2016). The distribution patterns of pathogens, parasites and their vectors are strongly dependent on temperature and humidity (Altizer et al. 2006) (Vajana et al. 2018), with for example a higher prevalence of the liver fluke under wet conditions or of several ectoparasites under hot and humid environments (Taylor 2012). In this context, selection signatures for genes involved in stem cells and immune B cell differentiation are likely to be related to environmental variation in pathogen selective pressures.

Another interesting case is that of the “Regulation of bone remodelling” category (GO:0046850), which is enriched in genes putatively selected in goats in association with solar radiation, namely *LRP5* and *CA2*. The latter, which is one of the six genes putatively selected in both sheep and goats, was also associated with solar radiation in sheep. We can therefore predict a possible role of these genes in preventing the decrease in bone density following lower levels of vitamin D at low UV radiation exposure. Indeed, vitamin D levels which affect bone density (Reid et al. 2014) depend on food intake and cutaneous production in sheep and goats (Kohler et al. 2013).

### Co-occurrence of anthropogenic and natural selection

Besides natural selection, anthropogenic selection can also have a strong impact on the genomes of domestic animals and disentangling both is not straightforward. In sheep, *MC5R* and *MC2R,* which are located in the largest selective sweeps identified (~650 kb), are related to temperature and rainfall. In *MC5R* we found a non-synonymous polymorphism spatially segregated in regions with the lowest rainfall and highest temperature. *MC5R* is expressed in a variety of peripheral tissues, especially in the skin (Switonski et al. 2013), and is a candidate gene for fat deposition in domestic animals because it down-regulates leptin secretion (Kristiansen and Mandrup-Poulsen 2005). Fat deposition, which is advantageous under dry conditions, could be directly selected by climatic pressures but also by anthropogenic selection in breeds such as the D’man, which is currently predominant in the dry and warm regions of Morocco. We also report several genes associated with rainfall, which may have been anthropogenically selected. They include *RXFP2* involved in horn morphology (Johnston et al. 2011) and reproductive success (Johnston et al. 2013), *RANBP17* associated to sperm maturation (Koch et al. 2000) (Bao et al. 2011), *SEMA5A* related to milk production traits and mammary gland inflammation in cattle (Sugimoto et al. 2006) (Pareek 2010), and *RBM19* involved in mammary gland morphology in cattle (Pausch et al. 2012). However, we currently cannot disentangle whether these observations are a combined effect of human-mediated and natural selection on the same traits, or the result of a confounding effect based on breeds adapted to a specific climate.

### Convergence/divergence of adaptive pathways

The extent to which genomic convergence shapes locally adapted phenotypes in different populations or species requires inferring the genetic bases of convergent phenotypes occurring in similar environments, see *e.g.,* (Wood et al. 2005) (Christin et al. 2010) (Elmer and Meyer 2011) (Conte et al. 2012). In this case, convergences at the genetic level would have been expected to be common as the probability of gene reuse is high and increases with closer common ancestry of the populations (Conte et al. 2012), such as for the two closely related species studied here. However, our results are in accordance with previous studies reporting that adaptation may involve convergence at different biological levels (from the same mutation to the production of different phenotypes for fulfilling the same organismal function) and that non-convergence at lower hierarchical levels may produce convergence at higher levels (Losos 2011) (Riesch et al. 2018). Out of the 420 putatively adaptive genes identified in this study, only six were orthologous between sheep and goats. Furthermore, the GO categories identified for the same environmental variable were always different. This supports the predominance of distinct genes and biological pathways for adaptation to the same environment in these species. There is growing evidence for the use of alternative genes or pathways in closely related species (Elmer and Meyer 2011) (Losos 2011) (Raeymaekers et al. 2017) or even in populations of the same species (Hoekstra and Nachman 2003) (Manceau et al. 2010). In addition, the global extent of gene reuse might be overestimated due to ascertainment and publication bias (Losos 2011). Less attention may also have been paid to convergent phenotypes resulting from different genes or pathways, and has been a bias towards the occurrence of gene reuse in studies based on candidate gene approaches (Gompel and Prud’homme 2009) (Conte et al. 2012). Whole genome approaches such as that used here provide a more realistic assessment of the extent of gene reuse in the determination of convergent phenotypes, which, though common, can mostly be considered as exceptions (Elmer and Meyer 2011).

Convergence at the genetic level is likely to result from constraints related to the size of mutational targets, the architecture of gene networks or the pleiotropic effect of impacted genes (Gompel and Prud’homme 2009) (Christin et al. 2010) (Losos 2011) (Tiffin and Ross-Ibarra 2014). In our study, the rarity of genetic convergence may be related to (i) relatively large mutational targets, the number of target genes and levels of standing genetic variation (Conte et al. 2012) and (ii) the diversity of gene networks related to the biological functions required to adapt to the combination of environmental parameters varying along climatic gradients. However, such adaptive processes are not free of constraints and in addition to the (few) genes found in common between sheep and goats, specific regulatory networks may play a central role in local adaptation. For example, the gene *MCM3* associated with altitude in sheep in our study has been shown to regulate *HIF* genes regulating haemoglobin concentration (Hubbi et al. 2011) (Semenza 2011). We did not find evidence for selection of *HIF* paralogs here, but they are widely reported to be under selection at high altitude (Simonson et al. 2010) (Yi et al. 2010) (Wei et al. 2016) (Song et al. 2016) (Gou et al. 2014). Other candidate genes involved in the *HIF* pathway have already been found under selection in sheep at higher altitudes than the present study (>3,300 m ASL; (Yang et al. 2016)). This would either illustrate the evolution of similar adaptations by selection acting at different levels in the same genetic network in different species or populations or by different selection pressures in apparently similar environments (Fig. 4). However, as our study did not characterize such phenotypes, it is not possible to discern the extent to which the adaptive mechanisms pursued in our study rely on convergent or different phenotypic traits.

## Conclusion

Our study provides an unusual example of analysing the diversity of recently differentiated adaptive mechanisms responding to environmental gradients. It provides novel insights to the debate over the extent of convergent evolution in shaping adaptation and thus diversification (Gould and Lewontin 1979) (Conway Morris 2003). We show empirically that the genetic basis of adaptive responses to the same climatic pressures are largely different in two closely related species, despite the high levels of genetic convergence between sheep and goats during/under the domestication process (Alberto et al. 2018). These findings are likely to result from the large mutational targets in these populations, which are related to the diversity of phenotypes, biological pathways and genes. The latter could be involved to increase fitness in the context of local adaptation to combinations of eco-climatic variations, while key changes related to domestication would target traits offering fewer options to selection, several of them relying on pleiotropic genes (Alberto et al. 2018).

## Methods

### Sampling

The sampling area (~400,000 km^2^; latitude range [28°-36°]; Fig. 1) covered the whole range of contrasting environments across Morocco. A sampling grid consisting of 162 cells of 0.5° of longitude and latitude was established and a maximum of three animals was sampled per flock for three different flocks per cell for each species. We collected the samples between January 2008 and March 2012 for the NextGen European project (KBBE-2009-1-1-03) in accordance with ethical regulations of the European Union Directive 86/609/EEC. For each individual, tissue samples were collected from the distal part of the ear and placed in alcohol for one day, and then transferred to a silica-gel tube until DNA extraction. A total of 412 flocks were sampled from which we aimed to select 164 individuals for Whole Genome Sequencing while optimizing sample selection to include the widest possible range of environmental conditions. The second criterion was to take the geographic location into account to maximise the spread of individuals across the region to ensure spatial representativeness for all environments. We therefore first performed a principal component analysis (PCA) on the 117 environmental variables extracted from the Climatic Research Unit (CRU) dataset (New et al. 2002). The PCA allowed us to maximise the ecological distance between the farms (separately for sheep and goats). Afterwards, we performed an Ascendent Hierarchical Classification (AHC) on the first 7 PCA-axes (96% of the variance) to group sampled farms according to their ecological distances. Using the Ward criterion, we reduced the number of classes to 164. After having grouped farms, we selected one individual per class. In order to guarantee spatial representativeness, we performed 50 random samplings and chose the one with the maximum index of distribution (i.e. the maximum sum of distances between each farm and its nearest neighbour). After sequencing, we removed seven individuals with low sequence quality, ending up with 161 goats and 160 sheep (see Fig. 1).

### Production of WGS data

DNA extraction was done using the Puregene Tissue Kit from Qiagen® following the manufacturer’s instructions and whole genome sequence data was produced and processed as described in Benjelloun et al (Benjelloun et al. 2015). 500ng of DNA was sheared to a 150-700 bp range using the Covaris® E210 (Covaris, Inc., USA). Sheared DNA was used for Illumina® library preparation with a semi-automated protocol. Briefly, end repair, A-tailing and Illumina® compatible adaptor (BiooScientific) ligation was performed using the SPRIWorks Library Preparation System and SPRI TE instrument (Beckmann Coulter), according to the manufacturer’s protocol. A 300-600 bp size selection was applied in order to recover most of fragments. DNA fragments were amplified in 12 cycles PCR using the Platinum Pfx Taq Polymerase Kit (Life® Technologies) and Illumina® adapter-specific primers. Libraries were purified with 0.8x AMPure XP beads (Beckmann Coulter). After library profile analysis using the Agilent 2100 Bioanalyzer (Agilent® Technologies, USA) and qPCR quantification, libraries were sequenced using 100 base-length read chemistry in paired-end flow cell on the Illumina HiSeq2000 (Illumina®, USA).

### WGS data processing

Illumina paired-end reads for sheep were mapped to the sheep reference genome (OAR v3.1, GenBank assembly GCA_000317765.1 (Jiang et al. 2014)) and those for goats were mapped to the goat reference genome (CHIR v1.0, GenBank assembly GCA_000317765.1 (Dong et al. 2013)) using BWA mem (Li and Durbin 2009). The BAM files produced were then sorted using Picard SortSam and improved using Picard Markduplicates (http://picard.sourceforge.net), GATK RealignerTargetCreator, GATK IndelRealigner (DePristo et al. 2011) and Samtools calmd (Li et al. 2009). Variant calling was carried out using three different algorithms: Samtools mpileup (Li et al. 2009), GATK UnifiedGenotyper (McKenna et al. 2010) and Freebayes (Garrison and Marth 2012).

There were two successive rounds of variant site filtering. Stage 1 merged calls together from the three algorithms, whilst filtering out the lowest-confidence calls. A variant site passed if it was called by at least two different calling algorithms with variant quality > 30. An alternate allele at a site passed if it was called by any one of the calling algorithms, and the genotype count > 0. Filtering stage 2 used Variant Quality Score Recalibration by GATK. First, we generated a training set of the highest-confidence variant sites where (i) the site is called by all three variant callers with variant quality > 100, (ii) the site is biallelic (Palti et al. 2015) the minor allele count is at least 3 while counting only samples with genotype quality > 30. The training set was used to build a Gaussian model using the GATK VariantRecalibrator tool using the following variant annotations from UnifiedGenotyper: QD, HaplotypeScore, MQRankSum, ReadPosRankSum, FS, DP, Inbreeding Coefficient. The Gaussian model was applied to the full data set, generating a VQSLOD (log odds ratio of being a true variant). Sites were filtered out if VQSLOD < cut-off value. The cutoff value was set for each population by the following: Minimum VQSLOD = {the median value of VQSLOD for training set variants} - 3 * {the median absolute deviation VQSLOD of training set variants}. Measures of the transition/transversion ratio of SNPs suggest that this chosen cut-off criterion gave the best balance between selectivity and sensitivity. Genotypes were improved and phased using Beagle 4 (Browning and Browning 2013), and then filtered out where the genotype probability calculated by Beagle was less than 0.95.

In order to detect orthologous signals of selection between sheep and goats, we performed a cross-alignment between the two reference genomes as described in Alberto et al. (2018). First, we used the pairwise alignment pipeline from the Ensembl release 69 code base (Flicek et al. 2012) to align the reference genomes of sheep (OARv3.1) and goat (CHIR1.0). This pipeline uses LastZ (Harris 2007) to align at the DNA level, followed by post-processing in which aligned blocks are chained together according to their location in both genomes. The LastZ pairwise alignment pipeline is run routinely by Ensembl for all supported species, but the goat was not included in Ensembl by the time of our analysis. We produced two different inter-specific alignments, using sheep as the reference genome and goat as non-reference and *vice versa.* This was done to avoid bias toward either species, given that genomic regions of the reference species are forced to map uniquely to a single locus of the non-reference species, whereas non-reference genomic regions are allowed to map to multiple locations of the reference species. We obtained for segments of chromosomes of one reference genome the coordinates on the non-reference genome. Finally, for the SNPs discovered in one genus, we used the whole genome alignment with the reference genome of the other genus to identify the corresponding positions.

### Genetic diversity and population structure

Neutral genomic variation was characterized to evaluate the level of genetic diversity present in Moroccan sheep and goats. The total number of variants and the number of variants within each species were calculated. The level of nucleotide diversity (*π*) was calculated in each species and averaged over all of the biallelic and fully diploid variants for which all individuals had a genotype called using Vcftools (Danecek et al. 2011). The observed frequency of heterozygous genotypes per individual (*Ho*) was calculated considering only the biallelic SNPs with no missing genotype calls. From *Ho,* inbreeding coefficients (*F*) were calculated for each individual using population allele frequencies over all individuals. F-tests were applied to test for the significance of differences between sheep and goat *Ho* and *F*.

Genetic structure was assessed using two different methods. First, we performed a principal component analysis (PCA) using an *LD* pruned subset of bi-allelic SNPs. *LD* between SNPs in windows containing 50 markers was calculated before removing one SNP from each pair where *LD* exceeded 0.95. Subsequently, only 12,543,534 SNPs among a total of 29,427,980 bi-allelic SNPs were kept for this analysis in goats and 14,056,772 out of 30,069,299 of biallelic SNPs for sheep. We used Plink v1.90a (https://www.cog-genomics.org/plink2) for *LD* pruning and the R package adegenet v1.3-1 (Jombart and Ahmed 2011) to run the PCA. Second, we carried-out an analysis with the clustering method sNMF (Frichot et al. 2014). This method was specifically developed for fast analysis of large genomic datasets. It is based on sparse non-negative matrix factorization to estimate admixture coefficients of individuals. We used all bi-allelic variants and performed five runs for each number of genetic clusters (i.e., K value varying from 1 to 10) using a value of *alpha* parameter of 16. For each run, we calculated the cross-entropy criterion with 5% missing data to identify the most likely number of clusters. We considered the run showing the lowest cross-entropy (CE) as the most likely value of K. Similarly we considered the number of clusters associated with the lowest CE as the most likely representative of our data structure.

### Environmental variables

We extracted the values of 67 climatic variables from the WorldClim dataset (Hijmans et al. 2005) (http://www.worldclim.org/current) for the sampling locations. These variables are based on data collected over 30 years and provide temperature and rainfall measurements as well as bioclimatic indices (i.e. derived from the monthly temperature and rainfall values in order to generate more biologically meaningful variables) with an initial resolution of 1×1 km. Additionally, we used a Digital Elevation Model (DEM) with a resolution of 90m (SRTM; http://earthexplorer.usgs.gov; courtesy of the U.S. Geological Survey) to obtain topography related variables. We computed a total of 4 DEM-derived variables with SAGA GIS (Böhner et al. 2006): altitude, slope and solar radiation in June and December. Afterwards, we conducted a pairwise correlation analysis between all 71 variables to detect highly correlated ones (|r|>=0.8), and kept 10 representative variables to investigate signatures of selection (Supplementary Tables 1 and 2).

### Analyses of signatures of selection

We applied two approaches to identify putative signatures of selection. The first was a correlative approach processing multiple parallel association models by means of logistic regressions between the frequencies at each locus with the value of the environmental variables selected at the 160 and 161 locations for sheep and goats, respectively. For this analysis, we used Samβada (Stucki et al. 2017), an individual-based method performing logistic regressions. This is a variant of linear regression in which the binary genetic marker is either present or absent and correlates with a quantitative environmental variable. Therefore, it provides the probability of occurrence of a genotype for each individual as a function of environmental gradients (Joost et al. 2007). It also provides spatial statistics that are helpful for the interpretation of significant results regarding spatial autocorrelation and is efficient in terms of computing time in the context of several million SNPs to process with 10 environmental variables. We considered only SNPs with a MAF>=0.05. Because Samβada requires binomial data, each biallelic SNP was coded in three columns corresponding to its genotypes. We then performed univariate regressions on each genotype with each of the 10 variables. Thereafter, the false discovery rate was calculated for each variable separately and Q-values were computed. The false discovery threshold was set to 0.1 on Samβada’s results to identify candidate SNPs.

The second approach was population-based and compared, through genome scans, two groups from contrasting conditions for each environmental variable. More precisely, for each variable, each of the two groups was constituted by the 20 individuals submitted to the most extreme values of the variable, at each side of the environmental gradient. Then, we ran XP-CLR (Chen et al. 2010) to identify potential genomic regions differentially selected between groups. This method is designed to identify regions under positive selection in an object population, given the other population considered as the reference. Therefore, we carried-out each analysis twice, each group being in turn the reference and the object population. XP-CLR is robust to detect selective sweeps and especially with regard to the uncertainty in the estimation of local recombination rate (Chen et al. 2010). Due to the absence of reliable information on genetic linkage for the whole genome in both species, the physical position (1 Mb ≈ 1 cM) was used. We used overlapped segments of a maximum of 27 Mb to estimate and assemble XP-CLR scores using the whole set of bi-allelic variants as described in Benjelloun et al. (Benjelloun et al. 2015). Segments were overlapping by 2Mb and the scores computed for the extremes 1Mb were discarded, except at the starting and end of chromosomes. XP-CLR scores were calculated using grid points spaced by 2500 bp with a minimum of 200 and a maximum of 250 variants in a window of 0.1 Mb, and by down-weighting contributions of highly correlated variants (*r*>0.95) in the reference group. The 0.01% genomic regions with the highest XP-CLR scores were identified. Within these regions (0.1 Mb each), the top differentiated variants between the two groups were defined applying a *Fst* (Weir and Cockerham 1984) cut-off level of 0.05% of the genome-wide *Fst* distribution. This threshold was chosen for having the highest enrichment values for outliers within XP-CLR windows (Student *t*-test, *p=7.3e-13* for sheep and *p=5.2e-13* for goats and *F*-test, *p=1.1e-37* for sheep and *p=1.6e-32* for goats). In addition, overlapping top XP-CLR regions were merged and all regions were ranked according to their XP-CLR score. Then, we checked for the occurrence of orthologous outlier variants, genes and genomic regions between sheep and goats. For that, outlier variants identified in one species were mapped on the genome assembly of the other species.

We used the Variant Effect Predictor (VEP) tools (McLaren et al. 2010) to classify the variants as intronic, exonic, synonymous, missense and intergenic. Genes including at least a candidate variant or located at less than 5 kb away from it (downstream 5’-end and upstream 3’-end) were used for Gene Ontology (GO) enrichment analyses. These analyses were done for each species using the sets of genes identified for each variable separately.

For each species we computed the profiles of allelic frequency variation for each selected region along the corresponding environmental gradient. For this analysis, we subdivided the range of values in 8 groups corresponding each to 20 contiguous individuals (except one group of 21 goats) for each environmental variable, and calculated the allelic frequency of the most differentiated candidate variant for each group (on the basis of its *Fst* value) associated with each genomic region. A neutral profile of allelic frequencies was represented by calculating for each case the allelic frequency of variants randomly selected over the genome (i.e., 41 for goats and 56 for sheep). We classified all profiles by comparing them to synthetic profiles corresponding to 3 different classes of pattern of variation: (i) a gradual linear variation of the frequency along the gradient, (ii) a uniform frequency all along the gradient except at one extreme end of the gradient and (Palti et al. 2015) a uniform frequency along the gradient except at the two extremes. We compared the frequency profiles with the synthetic ones using the Pearson correlation coefficient and attributed each profile to the synthetic profile exhibiting the highest absolute value for Pearson correlation.

### Gene Ontology enrichment analyses

To explore the biological processes involving the whole set of candidate genes identified by at least one method (i.e. Samßada and/or XP-CLR/*Fst*), Gene Ontology (GO) enrichment analyses were performed using GOrilla (Eden et al. 2009). The 12,669 and 14,620 genes associated with a GO term in goats and sheep, respectively, were used as background references. We assessed the significance for each individual GO-identifier with P-values corrected using a FDR q-value according to Benjamini and Hochberg’s method (Benjamini and Hochberg 1995). GO terms identified for each variable were clustered into homogenous groups using REVIGO by allowing medium similarity (0.7) (Supek et al. 2011). Low similarity among GO terms in a group was applied and the weight of each GO term was assessed by its p-value.

## Acknowledgements

This work was supported by the European Union 7th framework project NEXTGEN (Grant Agreement no. 244356, coordinated by P.T.), the FACCE ERA-NET Plus project CLIMGEN (grant ANR-14-JFAC-0002-01, coordinated by M.W.B.), the LabEx OSUG@2020 (Investissements d’avenir – ANR10LABX56), the Wellcome Trust [WT108749/Z/15/Z], and the European Molecular Biology Laboratory (I.S., L.Cl., P.F.). The SRTM V3.0 data product used in Fig 1 was retrieved from the online Data Pool, courtesy of the NASA Land Processes Distributed Active Archive Center (LP DAAC), USGS/Earth Resources Observation and Science (EROS) Center, Sioux Falls, South Dakota, https://lpdaac.usgs.gov/data_access/data_pool.

The authors are grateful to Abdelmajid Bechchari, Mustapha Ibnelbachyr, Mouad Chentouf, Mohammed BenBati, Rachid Hadria, Mouloud Laghmir, Lahbib Haounou, El Mekki Hafiani, El Mustapha Sekkour, Ali Dadouch, Abdelkader Lberji, Chérif Er-rouidi, Mohamed Ouatiq, Btissam Hilal and Mustapha Sadek who contributed to the field sampling. They also thank Ludovic Orlando for fruitful discussions.

## Accession numbers

The variant calls and genotype calls used in this paper are archived in the European Variation Archive with accession ID ERZ019290 for sheep and ID ERZ020631 For goats. The data are accessible at ftp://ftp.ebi.ac.uk/pub/databases/nextgen/

## Author contributions

The paper represents the joint efforts of several research groups, most of whom were involved in the NEXTGEN project (coordinated by P.T.). P.T., F.P., E.C. and R.N. designed the study. P.T. and F.P. supervised the joint work in NEXTGEN. B.B., A.S., A.C., M.W.B., S.J., R.N., P.A.-M., and P.F. supervised the work of their research group.

B.B. and A.C. collected the samples. A.A. and S.E. produced whole-genome sequences. S.J. developed the sampling design and supervised Geographic Information Systems. I.S., L.Cl., E.C., S.E., F.Bo. and B.B. contributed to bioinformatic analyses. B.B., K.L., S.S., B.S., F.J.A., F.Bo., P.L., L.Co., F.Bi., F.P. and M.B. did the analyses. B.B., K.L., S.J., F.Bo. and F.P. produced the figures. B.B., K.L. and F.P. wrote the text with input from all authors and especially F. Bo, S.J. and P.O.-t.W

