## Supplementary text and tables for "Multiple adaptive solutions to face climatic constraints: novel insights in the debate over the role of convergence in local adaptation"

2- Univ. Grenoble Alpes, Univ Savoie Mont Blanc, CNRS, LECA, F-38000 Grenoble, France.

3- Laboratory of Geographic Information Systems (LASIG), School of Architecture, Civil and Environmental Engineering (ENAC), Ecole Polytechnique Fédérale de Lausanne (EPFL), Lausanne, Switzerland.

4- European Molecular Biology Laboratory, European Bioinformatics Institute, Wellcome Genome Campus, Hinxton, Cambridge, CB10 1SD, United Kingdom.

5 - School of Biosciences, Cardiff University, Museum Avenue, CF10 3AX, Wales, UK. Sustainable Places Research Institute, Cardiff University, 33 Park Place, Cardiff, UK.

6- GenPhySE, Université de Toulouse, INRAE, INPT, ENVT, Castanet-Tolosan, France.

7- PTP Science Park, Bioinformatics Unit, Via Einstein - Loc. Cascina Codazza, Lodi, Italy.

8- Génomique Métabolique, Genoscope, Institut François Jacob, CEA, CNRS, Univ. Evry, Université Paris-Saclay, 91057 Evry, France.

9- Genoscope, Institut de biologie François-Jacob, Commissariat à l'Energie Atomique CEA, Université Paris-Saclay, Evry, France.

10- Dipartimento di Scienze Animali, della Nutrizione e degli Alimenti, Facoltà di Scienze Agrarie, Alimentari e Ambientali, Università Cattolica del S. Cuore, via Emilia Parmense n. 84, 29122, Piacenza (PC), Italy.

11- BioDNA - Centro di Ricerca sulla Biodiversità e sul DNA Antico, Facoltà di Scienze Agrarie, Alimentari e Ambientali, Università Cattolica del S. Cuore, via Emilia Parmense n. 84, 29122, Piacenza (PC), Italy.

12- AIA Associazione Italiana Allevatori, 00161 Roma, Italy.

### Supplementary methods

- **Estimation of linkage disequilibrium**

Pairwise linkage disequilibrium (LD) between polymorphic sites was assessed in the studied genomic regions through the correlation coefficient ($r^2$). It was estimated in 5 segments of 2Mb on different chromosomes (physical positions between 5 Mb and 7 Mb on chromosomes 6, 11, 16, 21 and 26). LD was estimated either by using the whole set of reliable variants or after discarding rare variants with a minor allele frequency (MAF) less than 0.05. For both estimations, $r^2$ values between all pairs of bi-allelic variants (SNPs and indels) on the same segment were calculated using Vcftools. Inter-SNP distances (kb) were binned into the following 7 classes: 0–0.2, 0.2–1, 1–2, 2–10, 10-30, 30-60 and 60-120 kb and observed pairwise *LD* was averaged for each inter-SNP distance class and used to draw *LD* decay.

### Supplementary results and discussion

- **Linkage disequilibrium and genetic diversity**

Using the whole set of reliable variants, the genomic distance at which linkage disequilibrium decayed to less than 0.15 was 655 bp in sheep and 166 bp in goats. Moreover, $r^2$ decayed to less than 0.1 in 3.12 kb and 2.1 kb in sheep and goats respectively. When withdrawing rare variants (MAF<0.05), the average $r^2$ decayed to less than 0.2 in 3.6 kb in sheep and 5.8 kb in goats. It decayed to less than 0.15 in 4.4 kb in sheep and 8.1 kb in goats. Thus, *LD* values were highly influenced by rare variants and their removal reversed the ranking of LD values that became lower for sheep. This fact could also partly explain differences in heterozygosity and inbreeding coefficients between the two species. The *LD* values reported here are lower than those reported on other domestic animals (i.e. horses, cattle, pigs) where it largely exceeds 10 kb for $r^2$=0.20 (Villa-Angulo et al., 2009); (Wade et al., 2009); (McCue et al., 2012); (Ai, Huang, & Ren, 2013);

(Veroneze et al., 2013). These low values would mainly result from high effective population sizes and very common extensive breeding systems favouring high gene flows among Moroccan sheep (respectively goats) (Benjelloun et al., 2015).

- **Bases of adaptations**

The identification of GO categories enriched in candidate genes under selection might shed light on the metabolic pathways involved in the adaptive processes.

Interestingly, the GO categories related to altitude in sheep are associated with lung epithelial cells and ciliated cells differentiation and proliferation. This is consistent with the function of these cells, which are involved in gas exchange and mucus production (Mercer et al. 2006). Some types of such cells were shown to be numerous and prominent in llamas living at high altitude (Heath, Smith, & Harris, 1976) and to proliferate after exposure to hypoxia in rabbits (Uhlik, Konradova, Vajner, & Adaskova, 2005). The enrichment of these GO categories relies on the gene *NFIB*, which is essential for lung maturation in mice (Steele-Perkins et al., 2005) and would play a considerable role in gas exchange of highlander sheep..

Another candidate gene related to altitude is *OXR1*. This gene conserved among eukaryotes is protective against oxidative stress (Oliver et al., 2011) and could play a role in the adaptation to the variation of the oxygen level with altitude. The role of other genes is less straightforward. For example, a strong signal of selection is associated with *GMDS* (71 intronic variants and 8 downstream), which contributes to the biosynthesis of GDP-(L)-fucose which metabolism defect has been associated with leukocyte adhesion deficiency type II in humans (Karsan et al., 1998).We could speculate a possible involvement in the immune response linked to a pathogenic context covarying with altitude in the study area.

In several cases, the pathways involving genes under selection are only related to very general processes, which make it difficult to precisely depict the bases of adaptation. For example, in goats, GO terms enriched in selected genes associated to temperature were linked to insulin receptor signalling pathway and regulation of transport and ATP metabolic process (Supplementary Table 6). Even knowing that ion transport is temperature dependent (Landeira-Fernandez, Castilho, & Block, 2012) and involved in cold adaptation in mammals (Stevens & Kido, 1974), adaptive pathways cannot be inferred yet. Likewise, in sheep, enriched GO categories for several temperature-linked variables were related to cyclic adenosine monophosphate (cAMP) (Supplementary Table 5), a second messenger involved in many cellular processes in response to neurotransmitters and hormones (e.g. (Kawasaki et al., 1998); (de Rooij et al., 1998), (Ludwig, Margalit, Eismann, Lancet, & Kaupp, 1990)). However, the known role of some genes involved in these pathways (e.g., MC5R or RXFP2)  may give first insights on the traits under selection (see Main text).

**Supplementary table 1:** Climatic variables used in this study

| Variable | Meaning and unit |
|---|---|
| **prec1** | Average Monthly Precipitation (mm) – January |
| **prec2** | Average Monthly Precipitation (mm) – February |
| **prec3** | Average Monthly Precipitation (mm) – March |
| **prec4** | Average Monthly Precipitation (mm) – April |
| **prec5** | Average Monthly Precipitation (mm) – May |
| **prec6** | Average Monthly Precipitation (mm) – June |
| **prec7** | Average Monthly Precipitation (mm) – July |
| **prec8** | Average Monthly Precipitation (mm) – August |
| **prec9** | Average Monthly Precipitation (mm) – September |
| **prec10** | Average Monthly Precipitation (mm) – October |
| **prec11** | Average Monthly Precipitation (mm) – November |
| **prec12** | Average Monthly Precipitation (mm) – December |
| **tmax1** | Average Monthly Maximum Temperature (°C *10) – January |
| **tmax2** | Average Monthly Maximum Temperature (°C *10) – February |
| **tmax3** | Average Monthly Maximum Temperature (°C *10) – March |
| **tmax4** | Average Monthly Maximum Temperature (°C *10) – April |
| **tmax5** | Average Monthly Maximum Temperature (°C *10) – May |
| **tmax6** | Average Monthly Maximum Temperature (°C *10) – June |
| **tmax7** | Average Monthly Maximum Temperature (°C *10) – July |
| **tmax8** | Average Monthly Maximum Temperature (°C *10) – August |
| **tmax9** | Average Monthly Maximum Temperature (°C *10) – September |
| **tmax10** | Average Monthly Maximum Temperature (°C *10) – October |
| **tmax11** | Average Monthly Maximum Temperature (°C *10) – November |
| **tmax12** | Average Monthly Maximum Temperature (°C *10) – December |
| **tmin1** | Average Monthly Minimum Temperature (°C *10) – January |
| **tmin2** | Average Monthly Minimum Temperature (°C *10) – February |
| **tmin3** | Average Monthly Minimum Temperature (°C *10) – March |
| **tmin4** | Average Monthly Minimum Temperature (°C *10) – April |
| **tmin5** | Average Monthly Minimum Temperature (°C *10) – May |
| **tmin6** | Average Monthly Minimum Temperature (°C *10) – June |
| **tmin7** | Average Monthly Minimum Temperature (°C *10) – July |
| **tmin8** | Average Monthly Minimum Temperature (°C *10) – August |
| **tmin9** | Average Monthly Minimum Temperature (°C *10) – September |
| **tmin10** | Average Monthly Minimum Temperature (°C *10) – October |
| **tmin11** | Average Monthly Minimum Temperature (°C *10) – November |
| **tmin12** | Average Monthly Minimum Temperature (°C *10) – December |

| | |
|---|---|
| **tmean1** | Average Monthly Mean Temperature (°C *10) – January |
| **tmean2** | Average Monthly Mean Temperature (°C *10) – February |
| **tmean3** | Average Monthly Mean Temperature (°C *10) – March |
| **tmean4** | Average Monthly Mean Temperature (°C *10) – April |
| **tmean5** | Average Monthly Mean Temperature (°C *10) – May |
| **tmean6** | Average Monthly Mean Temperature (°C *10) – June |
| **tmean7** | Average Monthly Mean Temperature (°C *10) – July |
| **tmean8** | Average Monthly Mean Temperature (°C *10) – August |
| **tmean9** | Average Monthly Mean Temperature (°C *10) – September |
| **tmean10** | Average Monthly Mean Temperature (°C *10) – October |
| **tmean11** | Average Monthly Mean Temperature (°C *10) – November |
| **tmean12** | Average Monthly Mean Temperature (°C *10) – December |
| **bio1** | Annual Mean Temperature |
| **bio2** | Mean Diurnal Range (Mean of monthly (max temp - min temp)) |
| **bio3** | Isothermality (BIO2/BIO7)(*100) |
| **bio4** | Temperature Seasonality (standard deviation *100) |
| **bio5** | Max Temperature of Warmest Month |
| **bio6** | Min Temperature of Coldest Month |
| **bio7** | Temperature Annual Range (BIO5-BIO6) |
| **bio8** | Mean Temperature of Wettest Quarter |
| **bio9** | Mean Temperature of Driest Quarter |
| **bio10** | Mean Temperature of Warmest Quarter |
| **bio11** | Mean Temperature of Coldest Quarter |
| **bio12** | Annual Precipitation |
| **bio13** | Precipitation of Wettest Month |
| **bio14** | Precipitation of Driest Month |
| **bio15** | Precipitation Seasonality (Coefficient of Variation) |
| **bio16** | Precipitation of Wettest Quarter |
| **bio17** | Precipitation of Driest Quarter |
| **bio18** | Precipitation of Warmest Quarter |
| **bio19** | Precipitation of Col |

**Supplementary Table 2:** Environmental variables correlated (|r|>=0.8) with the 10 variables retained for the analyses.

| Species | Retained variable | Correlated variables \|Pearson's r\|>=0.8 (Not retained) | | | | | | | | | | | | | | | |
|---|---|---|---|---|---|---|---|---|---|---|---|---|---|---|---|---|---|
| Sheep | Altitude | tmin_12 | tmin_11 | tmin_10 | tmin_9 | tmin_4 | tmin_3 | tmin_2 | tmin_1 | tmean_12 | tmean_11 | tmean_10 | tmean_3 | tmean_2 | tmean_1 | tmax_12 | Ti2106 |
| | Slope | | | | | | | | | | | | | | | | |
| | Ti2112 | | | | | | | | | | | | | | | | |
| | tmin_8 | tmin_9 | tmin_7 | tmin_6 | tmin_5 | tmin_4 | tmean_10 | tmean_9 | tmean_8 | tmean_7 | tmean_6 | tmean_5 | tmean_4 | tmean_3 | tmax_11 | tmax_10 | bio_1 |
| | tmean_7 | tmin_8 | tmin_7 | tmean_9 | tmean_8 | tmean_6 | tmean_5 | tmax_9 | tmax_8 | tmax_7 | tmax_6 | tmax_5 | tmax_4 | bio_10 | bio_5 | | |
| | prec_4 | prec_12 | prec_11 | prec_10 | prec_6 | prec_5 | prec_3 | prec_2 | prec_1 | bio_19 | bio_16 | bio_13 | bio_12 | | | | |
| | bio_15 | prec_9 | prec_8 | prec_7 | | | | | | | | | | | | | |
| | bio_8 | | | | | | | | | | | | | | | | |
| | bio_7 | tmax_7 | bio_5 | bio_4 | bio_2 | | | | | | | | | | | | |
| | bio_3 | | | | | | | | | | | | | | | | |
| Goats | Altitude | tmin_12 | tmin_11 | tmin_10 | tmin_4 | tmin_3 | tmin_2 | tmin_1 | tmean_12 | tmean_11 | tmean_2 | tmean_1 | tmax_12 | bio_11 | bio_6 | | |
| | Slope | | | | | | | | | | | | | | | | |
| | Ti2112 | | | | | | | | | | | | | | | | |
| | tmin_8 | tmin_9 | tmin_7 | tmin_6 | tmin_5 | tmin_4 | tmean_10 | tmean_9 | tmean_8 | tmean_7 | tmean_6 | tmean_5 | tmean_4 | tmean_3 | tmax_11 | tmax_10 | bio_1 |
| | tmean_7 | tmin_8 | tmin_7 | tmean_9 | tmean_8 | tmean_6 | tmean_5 | tmax_9 | tmax_8 | tmax_7 | tmax_6 | tmax_5 | tmax_4 | bio_10 | bio_5 | | |
| | prec_4 | prec_11 | prec_10 | prec_6 | prec_5 | prec_3 | prec_2 | prec_1 | bio_19 | bio_16 | bio_13 | bio_12 | | | | | |
| | bio_15 | prec_9 | prec_8 | prec_7 | | | | | | | | | | | | | |
| | bio_8 | | | | | | | | | | | | | | | | |
| | bio_7 | tmax_8 | tmax_7 | bio_5 | bio_4 | bio_2 | | | | | | | | | | | |
| | bio_3 | | | | | | | | | | | | | | | | |

All variables were not kept for the analyses because of their high correlation. prec, tmin, tmax and tmean are rainfall, lower temperature, higher temperature and mean temperature respectively for each month specified from 1 to 12. Variables starting with “bio” are bioclimatic variables derived from temperature and rainfall and presented at http://worldclim.org/bioclim.

**Supplementary Table 3:** Candidate genes associated with environmental parameters

| Environmental parameter | Altitude | | Slope | | Solar radiations | | Temperature | | | |
|---|---|---|---|---|---|---|---|---|---|---|
| WorldClim and DEM derived Variables | Altitude | | Slope | | Ti2112 | | Tmin8 | | Tmean7 | |
| Species | Sheep | Goats | Sheep | Goats | Sheep | Goats | Sheep | Goats | Sheep | Goats |
| # genomic regions | 22 | 62 | 19 | 22 | 23 | 35 | 29 | 29 | 38 | 24 |
| Candidate genes by population-based approach | **Gene pool1**<br>GMDS<br>MCM3<br>OXR1<br>ZNF831<br>ENSOARG00000021651<br><br>**Gene pool2**<br>NFIB<br>GMDS<br>U6<br>IRAK4<br>PUS7L<br>TWF1<br>TRPM3<br>ANKRD13D<br>SSH3<br>TPM1 | **Gene pool1**<br>KHDRBS2<br>DCLK1<br>PRAMEF12<br>KCNH3<br>SPATS2<br>**DENND4C**<br>RPS6<br>DST<br><br>**Gene pool2**<br>LOC102180242<br>PMPCA<br>SDCCAG3<br>SNAPC4<br>KCNH3<br>SPATS2<br>HPSE2<br>CCDC85A<br>**FAT3**<br>RUNX1<br>PXK<br>RPP14<br>CAMTA1<br>DCLK1<br>CCDC159<br>EPOR<br>LOC102189026<br>RAB3D<br>RGL3<br>SWSAP1<br>EXT1<br>DOCK10<br>FAM126A<br>PRAMEF12<br>ALDH4A1<br>PNPLA5 | **Gene pool1**<br>ENSOARG00000003995<br>NEUROG3<br>SV2B<br>DNAH3<br>ENSOARG00000014365<br>PAPD5<br><br>**Gene pool2**<br>ENSOARG00000020977<br>ENSOARG00000000908 | **Gene pool1**<br>C14H8orf37<br>ACD<br>AGRP<br>ATP6V0D1<br>C18H16orf86<br>CTCF<br>ENKD1<br>FAM65A<br>GFOD2<br>PARD6A<br>RLTPR<br>SNTG2<br>**DUSP2**<br>ACTR3B<br>KCNK6<br>MTF1<br><br>**Gene pool2**<br>PHTF1<br>EXD2<br>ANAPC16 | **Gene pool1**<br>**CA2**<br>5s_rRNA<br>CAB39L<br>AGR2<br>TSPAN13<br>CSNK1D<br><br>**Gene pool2**<br>CSRP2BP<br>OVOL2<br>PET117<br>DENND5B<br>ENSOARG00000008472<br>U6<br>ZNF140<br>ZNF26<br>ZNF268<br>SUPT3H<br>RORA<br>IQCA1<br>PIK3C3<br>ANO4<br>KRT71<br>FAM210A<br>LDLRAD4<br>HERC3<br>NAP1L5 | **Gene pool1**<br>LRP5<br>SAPS3<br>ANKS1B<br>SLC1A7<br><br>**Gene pool2**<br>CNOT7<br>ZDHHC2<br>LOC102180852<br>LOC102183418<br>FAM217B<br>PHACTR3<br>PPP1R3D<br>SYCP2<br>TBC1D1<br>PLIN3<br>TICAM1<br>HECW1<br>SAPS3<br>**CA2**<br>GUCY1A2<br>ATG4B<br>LOC102190576 | **Gene pool1**<br>ENSOARG00000016955<br>ABCG2<br>PKD2<br>BCAS1<br>DENND5B<br>OSBPL11<br>U6<br>ENSOARG00000013454<br>C22orf23<br>MICALL1<br>POLR2F<br>SNORA42<br>SOX10<br>ENSOARG00000009355<br>NT5DC1<br>TSPYL4<br>ENSOARG00000011042<br>MMP19<br>DNAH9<br>ENSOARG00000023834<br>FAM210A<br>LDLRAD4<br>MC5R<br>RNMT<br>DCAF10<br>ENSOARG00000011374<br>MC2R<br><br>**Gene pool2**<br>MC2R<br>MC5R<br>RNMT<br>ZNF831<br>ENSOARG00000008697<br>ENSOARG00000008723<br>PLD1<br>CRY1<br>ENSOARG00000017803<br>MTERF2<br>TMEM263 | **Gene pool1**<br>SPAG17<br>CATSPER3<br>SPG21<br>LOC102176945<br>LOC102180242<br>CCDC129<br>LMBRD1<br>STRIP1<br><br>**Gene pool2**<br>OXSR1<br>ESD<br>HTR2A<br>GRM7<br>HS6ST3<br>CDK6<br>PID1<br>KIAA0368 | **Gene pool1**<br>ENSOARG00000011616<br>RXFP2<br>ENSOARG00000001878<br>STK19<br>IMPA1<br>SLC10A5<br>ZFAND1<br>FAM210A<br>LDLRAD4<br>MC2R<br>MC5R<br>RNMT<br>ENSOARG00000011042<br>MMP19<br>OTOA<br>BCAS1<br><br>**Gene pool2**<br>ENSOARG00000015390<br>ENSOARG00000015647<br>ENSOARG00000011616<br>RXFP2<br>MC2R<br>MC5R<br>RAB35<br>ENSOARG00000011119<br>ENSOARG00000011124<br>ENSOARG00000017428<br>OSBPL1A<br>SNORA11<br>USP40<br>7SK<br>ENSOARG00000002492 | **Gene pool1**<br>SPAG17<br>MTDH<br>DEGS2<br>SAMD4A<br>ABCA4<br>SNRPC<br>UHRF1BP1<br>CYSTM1<br>PFDN1<br>KDM5B<br>ZNF536<br>LOC102181667<br><br>**Gene pool2**<br>TBCK<br>PET112<br>LOC102181593<br>BARD1<br>WRAP73 |

| | | | | | | | | | |
|---|---|---|---|---|---|---|---|---|---|
| | | | | | | | *ENSOARG00000015005*<br>*IFI30*<br>*IL12RB1*<br>*U4* | | |
| Candidate genes by correlative approach | | *PXK*<br>*GRK5*<br>*ARHGAP22*<br>*FKBP14*<br>*PLEKHA8*<br>*GRM1* | | | | | | | *MC5R*<br>*RNMT*<br>*USP40*<br>*SNORA11*<br>*ENSOARG00000011616*<br>*MC2R*<br>*LDLRAD4*<br>*TTC39C*<br>*TSPAN9*<br>*TMC6*<br>*TMPRSS15*<br>*FAM210A*<br>*DTYMK* |

| Environmental parameter | Rainfall | | Temperature seasonality | | | | | | Rainfall seasonality | |
|---|---|---|---|---|---|---|---|---|---|---|
| WorldClim and DEM derived Variables | Prec4 | | Bio7 | | Bio3 | | Bio8 | | Bio15 | |
| Species | Sheep | Goats | Sheep | Goats | Sheep | Goats | Sheep | Goats | Sheep | Goats |
| # genomic regions | 86 | 32 | 51 | 58 | 30 | 37 | 24 | 30 | 14 | 54 |

| | | | | | | | | | | |
|---|---|---|---|---|---|---|---|---|---|---|
| Candidate genes by population-based approach | **Gene pool1**<br>*MC2R*<br>*MC5R*<br>*RNMT*<br>*ENSOARG00000011119*<br>*ENSOARG00000011124*<br>*ENSOARG00000017428*<br>*OVOL2*<br>*SUPT3H*<br>*ENSOARG00000016183*<br>*ENSOARG00000015005*<br>*ENSOARG00000003965*<br>*ENSOARG00000002284*<br>*LRRC41*<br>*LURAP1*<br>*RAD54L*<br><br>**Gene pool2**<br>*FAM210A*<br>*LDLRAD4*<br>*MC2R*<br>*MC5R*<br>*RNMT*<br>*ENSOARG00000016955*<br>*ENSOARG00000003965*<br>*ENSOARG00000012123*<br>*MTCL1*<br>*ENSOARG00000004066*<br>*DNAH9*<br>*ENSOARG00000023834* | **Gene pool1**<br>*C16H1orf53*<br>*LHX9*<br>*DUSP6*<br>***BICC1***<br>*CDK6*<br><br>**Gene pool2**<br>*CDAN1*<br>*STARD9*<br>*TTBK2*<br>*ADAM28*<br>*ADAM7*<br>*ADAMDEC1*<br>*RECK*<br>*LOC102171608*<br>*ZBTB16*<br>*HAUS2*<br>*LRRC57*<br>*KCNG1* | **Gene pool1**<br>*GMDS*<br>*ENSOARG00000023809*<br>*FOXP4*<br>*RYBP*<br>***TCP11L2***<br>*SALL3*<br>*ENSOARG00000022011*<br>*TBX19*<br>*AGMO*<br>*ENSOARG00000012660*<br>*CHN2*<br>*MAP3K11*<br>*PCNXL3*<br>*ENSOARG00000004627*<br>*ENSOARG00000004657*<br>*U6*<br>*CNNM2*<br>*uc_338*<br><br>**Gene pool2**<br>*ENSOARG00000022526*<br>*LAMB1*<br>*ENSOARG00000008550*<br>*EDAR*<br>*PTPN4*<br>*ENSOARG00000023181*<br>*RAMP1*<br>*GMDS* | **Gene pool1**<br>*ADAMTS20*<br>*JARID2*<br>*KIFAP3*<br>*CASC4*<br><br>**Gene pool2**<br>*GLS*<br>*LOC102189170*<br>*LACTB2*<br>*TRAM1*<br>*PKD2L1*<br>*PTGR1*<br>*ABCA4*<br>*CP*<br>*LOC102172487*<br>*PASK*<br>*PPP1R7*<br>*TMEM163* | **Gene pool1**<br>*ENSOARG00000015906*<br>***DENND4C***<br>*P3H2*<br>*ENSOARG00000006683*<br>*ENSOARG00000006689*<br>*ZBTB42*<br>*FHOD3*<br>*ENSOARG00000015338*<br>*ENSOARG00000015398*<br>*MUC12*<br><br>**Gene pool2**<br>*ATP8B4*<br>*SLC27A2*<br>*ERMARD*<br>*PHF10*<br>*TCTE3*<br>*DCAF10*<br>*ENSOARG00000008225*<br>*ENSOARG00000011374*<br>*SHB*<br>*BCS1L*<br>*STK36*<br>*ZNF142*<br>*BTBD16*<br>*ENSOARG00000022395*<br>*TACC2*<br>*CCDC126*<br>*DBF4*<br>*ASTL*<br>***DUSP2***<br>*ENSOARG00000016583*<br>*STARD7*<br>*IRG1*<br>*ZNF605* | **Gene pool1**<br>*ZNF704*<br>*POLR3B*<br>***TCP11L2***<br>*ERLIN1*<br>*LOC102181835*<br>*PKD2L1*<br>*ARHGEF3*<br>*NRXN3*<br>*PLOD2*<br>*C6H4orf22*<br>*SRGAP3*<br>*SLC6A11*<br>*APRT*<br>*CDT1*<br>*GALNS*<br>*DYSF*<br>*NCOA7*<br>*RPTOR*<br><br>**Gene pool2**<br>*CMTM7*<br>*TG*<br>*KIAA1279*<br>*DHRS3*<br>*RIOK3*<br>*TMEM241*<br>*UCP2*<br>*UCP3* | **Gene pool1**<br>*7SK*<br>*CREB3L2*<br>*FAM210A*<br>*LDLRAD4*<br>*MC2R*<br>*MC5R*<br>*RNMT*<br>*HERC4*<br>*ENSOARG00000021087*<br>*ENSOARG00000000166*<br>*SND1*<br>*EME1*<br>*LRRC59*<br>*MRPL27*<br>*XYLT2*<br><br>**Gene pool2**<br>*MC2R*<br>*MC5R*<br>*STPG2*<br>*ENSOARG00000016183*<br>*ITGA6*<br>*ENSOARG00000014022*<br>*ENSOARG00000022000* | **Gene pool1**<br>*LOC102191021*<br>*LOC102191308*<br>*ABL1*<br>*LAMC3*<br>*PLCB4*<br>*LOC102176755*<br>*PKMYT1*<br>*LOC102180242*<br><br>**Gene pool2**<br>*GRIN2D*<br>*LOC102187192*<br>*PAFAH1B1*<br>*GPR107*<br>*HMCN2*<br>*NCS1*<br>*ALS2*<br>*MPP4*<br>*NOTCH1*<br>*ARHGAP20*<br>*LOC102181593* | **Gene pool1**<br>*H2AFY*<br><br>**Gene pool2**<br>*ADGRB1*<br>*TSNARE1*<br>*ENSOARG00000008039*<br>*NKTR*<br>*SS18L2*<br>*HACD2*<br>*PLEKHA7* | **Gene pool1**<br>*CNTN5*<br>*CDH2*<br>*LYRM9*<br>*CUL7*<br>*KLC4*<br>*KLHDC3*<br>*MEA1*<br>*RRP36*<br>*MS4A13*<br>*LOC102190926*<br>*ARHGEF38*<br>*LOC102186025*<br>*RAPGEF1*<br>*HECTD4*<br>*AKAP9*<br>*NUDT9*<br>*PPP1CB*<br>*SPDYA*<br>*TRMT61B*<br>*WDR43*<br><br>**Gene pool2**<br>*LOC102175300*<br>*INPP4A*<br>*SAPS3*<br>*ARHGEF38*<br>*LOC102177669*<br>*LOC102177939*<br>*WDR48* |
| Candidate genes by correlative approach | *MC5R*<br>*RNMT*<br>*ENSOARG00000011616*<br>*RBM19*<br>*RANBP17*<br>*SNRK*<br>*PIEZO2*<br>*MC2R*<br>*SEMA5A*<br>*PKHD1*<br>*LDLRAD4*<br>*MIP*<br>*GREB1*<br>*EXT2*<br>*CNBD1*<br>***BICC1***<br>*AATF*<br>*WDR62*<br>*TRAPPC9*<br>*TRAF3IP1*<br>*TMC6*<br>*THSD7B*<br>*SNORA3*<br>*SFMBT2*<br>*NGFR*<br>*LIN28B* | | *EXD3*<br>*MC2R*<br>*MC5R*<br>*RNMT*<br>*SNORA11*<br>*THAP8*<br>*USP40*<br>*VAC14*<br>*ENSOARG00000019858*<br>*ENSOARG00000011616* | *ABCC5*<br>*KLF12*<br>*RPS6KC1*<br>*PTPRG*<br>*DLGAP1*<br>*KRT78*<br>*NKAIN2*<br>*MANEA* | | | | | | *AGTR1*<br>*FAM188A*<br>*DIRC1*<br>*DSG4*<br>*CDH2*<br>*KCTD1*<br>*CEP192*<br>*LOC102183037*<br>*WRN*<br>*FRMPD2*<br>*KIAA0319L*<br>*EXOC4*<br>*PPFIA2*<br>*UNC5C* |

| | |
|---|---|
| | ***FAT3***<br>*FAM210A*<br>*FAM114A1*<br>*DPEP1*<br>*B4GALNT2*<br>*ALX4*<br>*ABCG1* |

Highlighted genes in green correspond to homologous candidate genes in both sheep and goats. For each environmental variable, candidate genes identified in high-level pool and respectively low-level pool of individuals using XP-CLR/Fst were ranked according to their XP-CLR scores and displayed in Gene pool1 and respectively Gene pool2. Underlined genes were identified in more than one analysis (population based in High-level, population based in low-level, correlative).

**Supplementary Table 4 :** Annotation of outlier variants

| Variant group | Sheep | | Goats | |
|---|---|---|---|---|
| | # | % | # | % |
| **Missense variant** | 20 | 0.2 | 1 | 0 |
| **Synonymous variant** | 47 | 0.4 | - | - |
| **Missense variant/splice region variant** | 1 | 0.0 | - | - |
| **3' UTR variant** | 54 | 0.5 | 54 | 1.0 |
| **5' UTR variant** | 7 | 0.1 | 11 | 0.2 |
| **Intron variant** | 3020 | 26.6 | 1335 | 24.0 |
| **Splice region variant/intron variant** | 7 | 0.1 | 4 | 0.1 |
| **Upstream gene variant** | 537 | 4.7 | 225 | 4.0 |
| **Downstream gene variant** | 563 | 5.0 | 169 | 3.0 |
| **Intergenic variant** | 7111 | 62.5 | 3857 | 68.2 |

**Supplementary Table 5:** Enrichment analysis for candidate genes in relation with environmental variables in sheep

| Environmental variable | GO Term | Biological process | P-value | Enrichment | Associated genes | Associated candidate genes | Genes |
|---|---|---|---|---|---|---|---|
| Altitude | GO:2000795 | negative regulation of epithelial cell proliferation involved in lung morphogenesis (a) | 0.00082 | 1219.58 | 1 | 1 | *NFIB* |
| | GO:0043538 | regulation of actin phosphorylation (b) | 0.00082 | 1219.58 | 1 | 1 | *TWF1* |
| | GO:0003065 | positive regulation of heart rate by epinephrine (c) | 0.00082 | 1219.58 | 1 | 1 | *TPM1* |
| | GO:0061141 | lung ciliated cell differentiation (a) | 0.00082 | 1219.58 | 1 | 1 | *NFIB* |
| Tmean7 | GO:0030819 | positive regulation of cAMP biosynthetic process (a) | 0.0000903 | 33.77 | 65 | 3 | *RXFP2, MC2R, MC5R* |
| | GO:0030816 | positive regulation of cAMP metabolic process (a) | 0.000118 | 30.92 | 71 | 3 | *RXFP2, MC2R, MC5R* |
| | GO:0030804 | positive regulation of cyclic nucleotide biosynthetic process (a) | 0.000174 | 27.1 | 81 | 3 | *RXFP2, MC2R, MC5R* |
| | GO:1900373 | positive regulation of purine nucleotide biosynthetic process (a) | 0.000208 | 25.53 | 86 | 3 | *RXFP2, MC2R, MC5R* |
| | GO:0030810 | positive regulation of nucleotide biosynthetic process (a) | 0.000208 | 25.53 | 86 | 3 | *RXFP2, MC2R, MC5R* |
| | GO:0030801 | positive regulation of cyclic nucleotide metabolic process (a) | 0.00023 | 24.67 | 89 | 3 | *RXFP2, MC2R, MC5R* |
| | GO:0030817 | regulation of cAMP biosynthetic process (a) | 0.000296 | 22.63 | 97 | 3 | *RXFP2, MC2R, MC5R* |
| | GO:0030814 | regulation of cAMP metabolic process (a) | 0.000407 | 20.33 | 108 | 3 | *RXFP2, MC2R, MC5R* |
| | GO:0045981 | positive regulation of nucleotide metabolic process (a) | 0.000441 | 19.78 | 111 | 3 | *RXFP2, MC2R, MC5R* |
| | GO:1900544 | positive regulation of purine nucleotide metabolic process (a) | 0.000441 | 19.78 | 111 | 3 | *RXFP2, MC2R, MC5R* |
| | GO:0030802 | regulation of cyclic nucleotide biosynthetic process (a) | 0.000489 | 19.09 | 115 | 3 | *RXFP2, MC2R, MC5R* |

| | | | | | | | |
|---|---|---|---|---|---|---|---|
| | GO:1900371 | regulation of purine nucleotide biosynthetic process (a) | 0.000567 | 18.14 | 121 | 3 | *RXFP2, MC2R, MC5R* |
| | GO:0030808 | regulation of nucleotide biosynthetic process (b) | 0.000581 | 17.99 | 122 | 3 | *RXFP2, MC2R, MC5R* |
| | GO:0030799 | regulation of cyclic nucleotide metabolic process (c) | 0.000731 | 16.63 | 132 | 3 | *RXFP2, MC2R, MC5R* |
| Bio7 | GO:0030819 | positive regulation of cAMP biosynthetic process (a) | 0.000105 | 32.16 | 65 | 3 | *RAMP1, MC2R, MC5R* |
| | GO:0030816 | positive regulation of cAMP metabolic process (a) | 0.000137 | 29.45 | 71 | 3 | *RAMP1, MC2R, MC5R* |
| | GO:0030804 | positive regulation of cyclic nucleotide biosynthetic process (a) | 0.000202 | 25.81 | 81 | 3 | *RAMP1, MC2R, MC5R* |
| | GO:1900373 | positive regulation of purine nucleotide biosynthetic process (a) | 0.000241 | 24.31 | 86 | 3 | *RAMP1, MC2R, MC5R* |
| | GO:0030810 | positive regulation of nucleotide biosynthetic process (a) | 0.000241 | 24.31 | 86 | 3 | *RAMP1, MC2R, MC5R* |
| | GO:0030801 | positive regulation of cyclic nucleotide metabolic process (a) | 0.000267 | 23.49 | 89 | 3 | *RAMP1, MC2R, MC5R* |
| | GO:0030817 | regulation of cAMP biosynthetic process (a) | 0.000344 | 21.55 | 97 | 3 | *RAMP1, MC2R, MC5R* |
| | GO:0030814 | regulation of cAMP metabolic process (a) | 0.000472 | 19.36 | 108 | 3 | *RAMP1, MC2R, MC5R* |
| | GO:0045981 | positive regulation of nucleotide metabolic process (a) | 0.000511 | 18.84 | 111 | 3 | *RAMP1, MC2R, MC5R* |
| | GO:1900544 | positive regulation of purine nucleotide metabolic process (a) | 0.000511 | 18.84 | 111 | 3 | *RAMP1, MC2R, MC5R* |
| | GO:0030802 | regulation of cyclic nucleotide biosynthetic process (a) | 0.000567 | 18.18 | 115 | 3 | *RAMP1, MC2R, MC5R* |
| | GO:1900371 | regulation of purine nucleotide biosynthetic process (a) | 0.000658 | 17.28 | 121 | 3 | *RAMP1, MC2R, MC5R* |
| | GO:0030808 | regulation of nucleotide biosynthetic process (b) | 0.000674 | 17.14 | 122 | 3 | *RAMP1, MC2R, MC5R* |
| | GO:0030799 | regulation of cyclic nucleotide metabolic process (c) | 0.000847 | 15.84 | 132 | 3 | *RAMP1, MC2R, MC5R* |

| | | | | | | | |
|---|---|---|---|---|---|---|---|
| Bio15 | GO:1904815 | negative regulation of protein localization to chromosome, telomeric region | 0.000683 | 1463.5 | 2 | 1 | *H2AFY* |
| Ti2112 | GO:0010719 | negative regulation of epithelial to mesenchymal transition | 0.000489 | 60.48 | 22 | 2 | *OVOL2, LDLRAD4* |
| Tmin8 | GO:0030814 | regulation of cAMP metabolic process | 0.000799 | 16.26 | 108 | 3 | *PKD2, MC2R, MC5R* |

For each environmental variable, biological processes marked with the same letter were clustered together using REVIGO (Supek et al. 2011) with medium similarity.

**Supplementary Table 6:** Enrichment analysis for candidate genes in relation with environmental parameters in goats.

| Environmental variable | GO Term | Biological process | P-value | Enrichment | Associated genes | Associated candidate genes | Genes |
|---|---|---|---|---|---|---|---|
| Bio3 | GO:0000303 | response to superoxide (a) | 0.0000499 | 180.86 | 7 | 2 | *UCP3, UCP2* |
| | GO:0000305 | response to oxygen radical (a) | 0.0000854 | 140.67 | 9 | 2 | *UCP3, UCP2* |
| | GO:0006839 | mitochondrial transport (b) | 0.000565 | 18.26 | 104 | 3 | *KIAA1279, UCP3, UCP2* |
| | GO:0032868 | response to insulin (a) | 0.000943 | 15.31 | 124 | 3 | *APRT, UCP3, UCP2* |
| Bio7 | GO:0043576 | regulation of respiratory gaseous exchange | 0.000302 | 76.63 | 20 | 2 | *PASK, GLS* |
| Bio8 | GO:0032989 | cellular component morphogenesis (a) | 0.00000702 | 16.67 | 336 | 5 | *PAFAH1B1, ABL1, LAMC3, NOTCH1, ALS2* |
| | GO:0050905 | neuromuscular process (b) | 0.0000225 | 52.5 | 64 | 3 | *PAFAH1B1 , ABL1, GRIN2D* |
| | GO:0048708 | astrocyte differentiation (a) | 0.0000768 | 149.32 | 15 | 2 | *ABL1, NOTCH1* |
| | GO:0007409 | Axonogenesis (a) | 0.0000961 | 32.31 | 104 | 3 | *PAFAH1B1, NOTCH1, ALS2* |
| | GO:0017145 | stem cell division (c) | 0.000168 | 101.81 | 22 | 2 | *PAFAH1B1 , NOTCH1* |
| | GO:1903053 | regulation of extracellular matrix organization (d) | 0.000275 | 79.99 | 28 | 2 | *ABL1, NOTCH1* |
| | GO:0007626 | locomotory behavior (e) | 0.000337 | 21.13 | 159 | 3 | *PAFAH1B1, GRIN2D, ALS2* |
| | GO:0048812 | neuron projection morphogenesis (a) | 0.000477 | 18.77 | 179 | 3 | *PAFAH1B1, NOTCH1, ALS2* |
| | GO:0050885 | neuromuscular process controlling balance (b) | 0.000482 | 60.54 | 37 | 2 | *PAFAH1B1, ABL1* |
| | GO:0048858 | cell projection morphogenesis (a) | 0.000534 | 18.06 | 186 | 3 | *PAFAH1B1, NOTCH1, ALS2* |
| | GO:0030514 | negative regulation of BMP signaling pathway (f) | 0.000651 | 52.09 | 43 | 2 | *ABL1, NOTCH1* |
| | GO:0032990 | cell part morphogenesis (a) | 0.000709 | 16.39 | 205 | 3 | *PAFAH1B1, NOTCH1, ALS2* |
| | GO:0045664 | regulation of neuron differentiation (a) | 0.000745 | 9.03 | 496 | 4 | *PAFAH1B1, ABL1, NOTCH1, NCS1* |
| | GO:0007067 | mitotic nuclear division (a) | 0.000848 | 15.41 | 218 | 3 | *PAFAH1B1 , ABL1, PKMYT1* |

| | | | | | | | |
|---|---|---|---|---|---|---|---|
| | GO:0002332 | transitional stage B cell differentiation (a) | 0.000893 | 1119.92 | 1 | 1 | *ABL1* |
| | GO:0002333 | transitional one stage B cell differentiation (a) | 0.000893 | 1119.92 | 1 | 1 | *ABL1* |
| | GO:0051882 | mitochondrial depolarization (g) | 0.000893 | 1119.92 | 1 | 1 | *ABL1* |
| | GO:1904618 | positive regulation of actin binding (a) | 0.000893 | 1119.92 | 1 | 1 | *ABL1* |
| | GO:1904531 | positive regulation of actin filament binding (a) | 0.000893 | 1119.92 | 1 | 1 | *ABL1* |
| | GO:0003213 | cardiac right atrium morphogenesis (a) | 0.000893 | 1119.92 | 1 | 1 | *NOTCH1* |
| | GO:0003270 | Notch signaling pathway involved in regulation of secondary heart field cardioblast proliferation (a) | 0.000893 | 1119.92 | 1 | 1 | *NOTCH1* |
| | GO:0060843 | venous endothelial cell differentiation (a) | 0.000893 | 1119.92 | 1 | 1 | *NOTCH1* |
| | GO:0090176 | microtubule cytoskeleton organization involved in establishment of planar polarity (a) | 0.000893 | 1119.92 | 1 | 1 | *PAFAH1B1* |
| | GO:0007440 | foregut morphogenesis (a) | 0.000893 | 1119.92 | 1 | 1 | *NOTCH1* |
| Bio15 | GO:0044772 | mitotic cell cycle phase transition (a) | 0.000615 | 10.04 | 223 | 4 | *AKAP9, PPP1CB, CEP192, SPDYA* |
| | GO:0044770 | cell cycle phase transition (a) | 0.000668 | 9.82 | 228 | 4 | *AKAP9, PPP1CB, CEP192, SPDYA* |
| Prec4 | GO:0035136 | forelimb morphogenesis | 0.000612 | 53.92 | 36 | 2 | *RECK, ZBTB16* |
| Slope | GO:0034502 | protein localization to chromosome | 0.000834 | 46.29 | 37 | 2 | *ACD, CTCF* |
| Ti2112 | GO:0046850 | regulation of bone remodeling | 0.000737 | 49.19 | 37 | 2 | *CA2, LRP5* |
| Tmin8 | GO:0051051 | negative regulation of transport (a) | 0.000018 | 13.9 | 374 | 5 | *GRM7, OXSR1, LMBRD1, HTR2A, PID1* |
| | GO:0046325 | negative regulation of glucose import (b) | 0.0000666 | 159.99 | 13 | 2 | *LMBRD1, PID1* |
| | GO:0043271 | negative regulation of ion transport (b) | 0.0000873 | 33.55 | 93 | 3 | *GRM7, OXSR1, HTR2A* |
| | GO:0010829 | negative regulation of glucose transport (b) | 0.000102 | 129.99 | 16 | 2 | *LMBRD1, PID1* |
| | GO:0045979 | positive regulation of nucleoside metabolic process (c) | 0.00013 | 115.55 | 18 | 2 | *HTR2A, PID1* |

| | | | | | | | |
|---|---|---|---|---|---|---|---|
| | GO:1903580 | positive regulation of ATP metabolic process (d) | 0.00013 | 115.55 | 18 | 2 | *HTR2A, PID1* |
| | GO:0043267 | negative regulation of potassium ion transport (b) | 0.000215 | 90.43 | 23 | 2 | *OXSR1, HTR2A* |
| | GO:0046627 | negative regulation of insulin receptor signaling pathway (e) | 0.00032 | 74.28 | 28 | 2 | *LMBRD1, PID1* |
| | GO:1900077 | negative regulation of cellular response to insulin stimulus (e) | 0.000368 | 69.33 | 30 | 2 | *LMBRD1, PID1* |
| | GO:0070588 | calcium ion transmembrane transport (f) | 0.000418 | 19.75 | 158 | 3 | *GRM7, HTR2A, CATSPER3* |
| | GO:1900372 | negative regulation of purine nucleotide biosynthetic process (d) | 0.000502 | 59.42 | 35 | 2 | *GRM7, PID1* |
| | GO:0030809 | negative regulation of nucleotide biosynthetic process (d) | 0.000502 | 59.42 | 35 | 2 | *GRM7, PID1* |
| | GO:1900542 | regulation of purine nucleotide metabolic process (d) | 0.000509 | 18.46 | 169 | 3 | *GRM7, HTR2A, PID1* |
| | GO:0006140 | regulation of nucleotide metabolic process (d) | 0.000554 | 17.93 | 174 | 3 | *GRM7, HTR2A, PID1* |
| | GO:1903578 | regulation of ATP metabolic process (d) | 0.000723 | 49.52 | 42 | 2 | *HTR2A, PID1* |
| | GO:0009118 | regulation of nucleoside metabolic process (g) | 0.000723 | 49.52 | 42 | 2 | *HTR2A, PID1* |
| | GO:0046626 | regulation of insulin receptor signaling pathway (e) | 0.000868 | 45.21 | 46 | 2 | *LMBRD1, PID1* |
| | GO:0006816 | calcium ion transport (f) | 0.000958 | 14.86 | 210 | 3 | *GRM7, HTR2A, CATSPER3* |
| | GO:0038016 | insulin receptor internalization (h) | 0.000962 | 1039.93 | 1 | 1 | *LMBRD1* |
| | GO:0007208 | phospholipase C-activating serotonin receptor signaling pathway (i) | 0.000962 | 1039.93 | 1 | 1 | *HTR2A* |
| | GO:0051966 | regulation of synaptic transmission, glutamatergic (j) | 0.000984 | 42.45 | 49 | 2 | *GRM7, HTR2A* |

For each environmental variable, biological processes marked with the same letter were clustered together using REVIGO (Supek et al. 2011) with medium similarity.
