## Supplementary figures for "Multiple adaptive solutions to face climatic constraints: novel insights in the debate over the role of convergence in local adaptation"

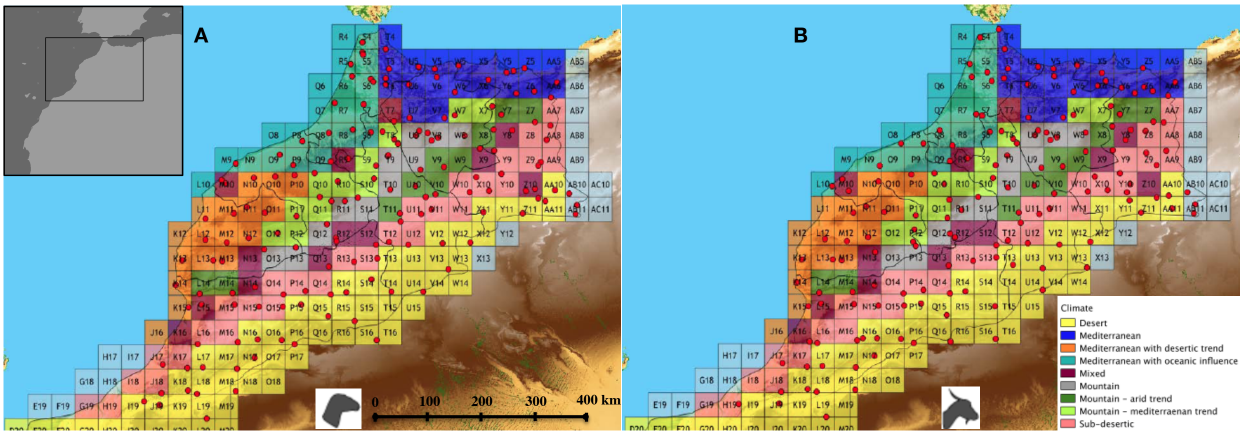


**Supp Fig 1**: Sampling grid with location of the sequenced individuals (A: sheep, B: goats) and the diversity of climate conditions over the study area in Morocco.

A

B

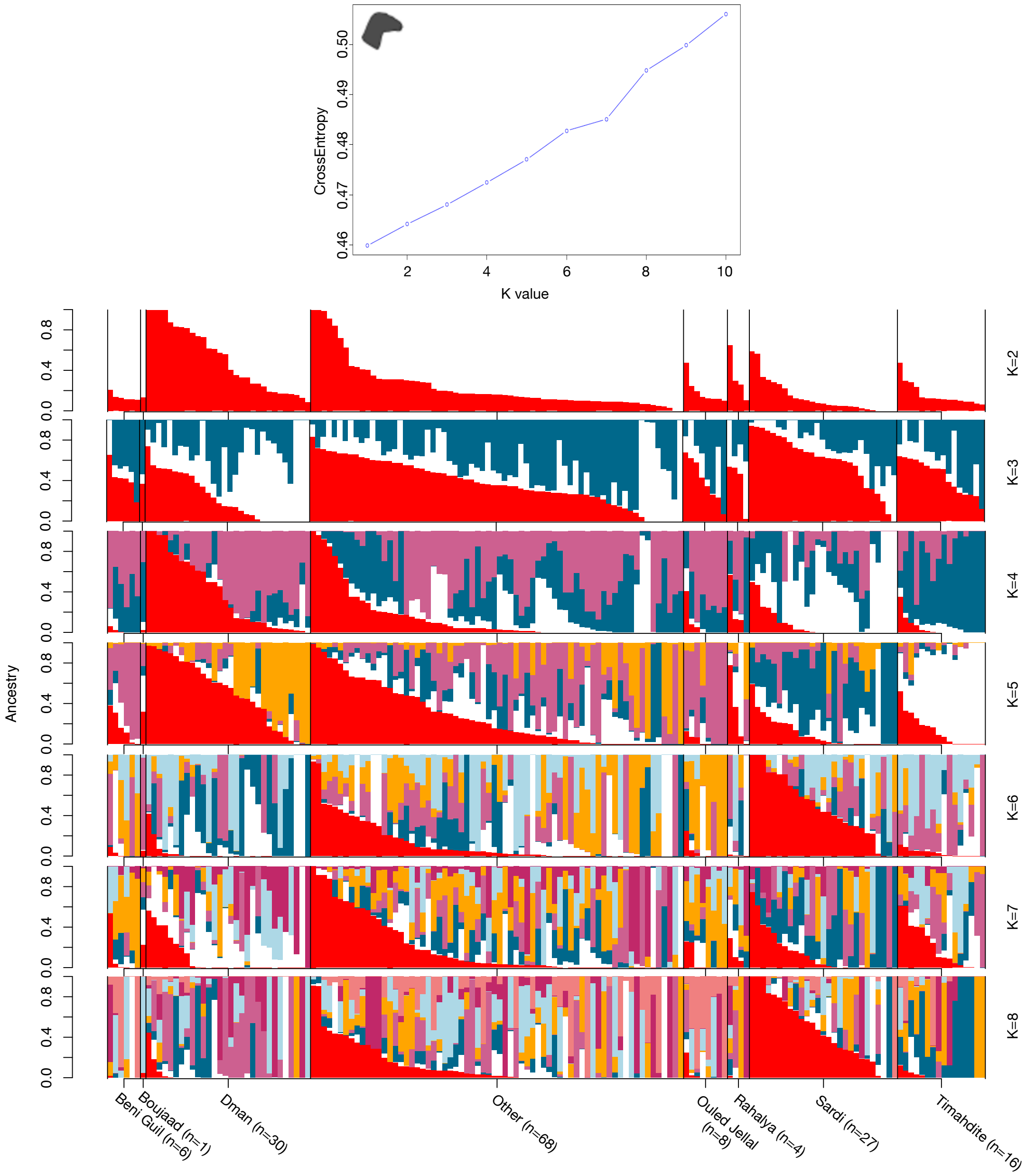


**Supp Fig 2**: sNMF analysis of whole genome data in sheep. A. Variation of cross entropy showing K=1 as the most likely number of genetic clusters. B. Proportions of genomes assigned to K genetic clusters (K=2 to 8), showing the absence of structure between breeds.

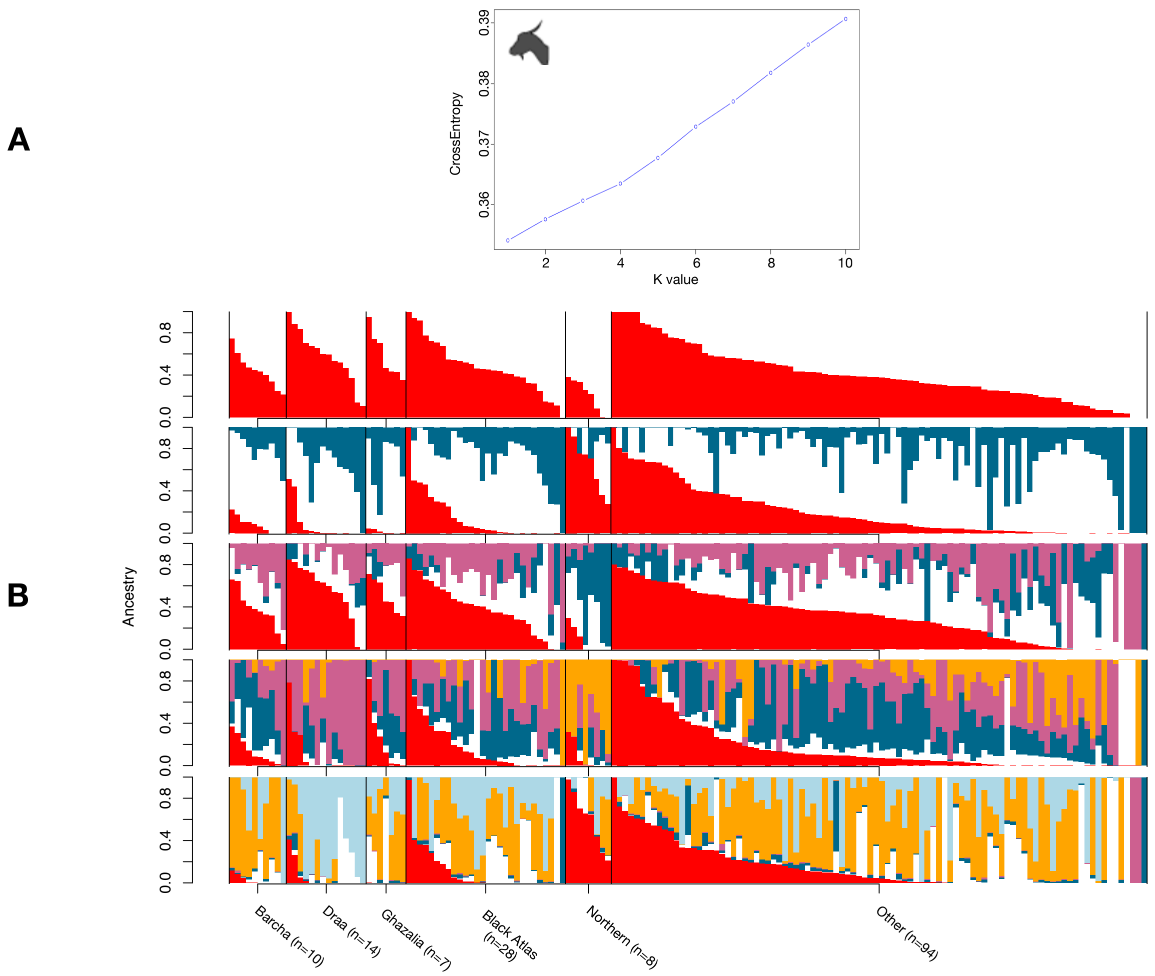


**Supp Fig 3**: sNMF analysis of whole genome data in goats. A. Variation of cross entropy showing K=1 as the most likely number of genetic clusters. B. Proportions of genomes assigned to K genetic clusters (K=2 to 6), showing the absence of structure between breeds.

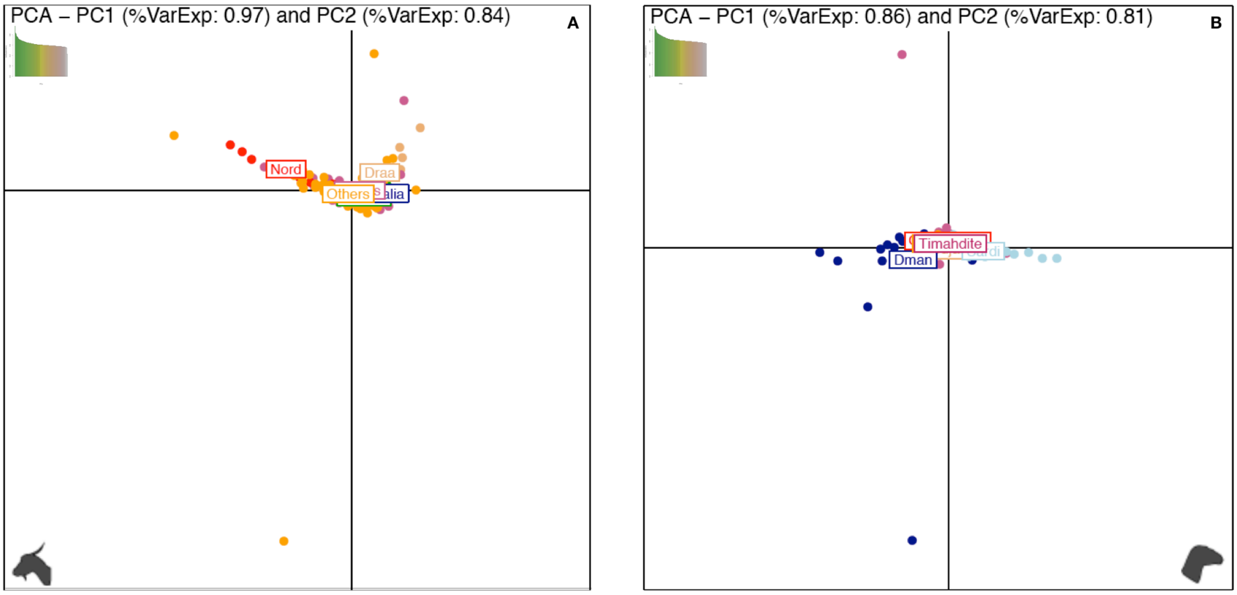


**Supp Fig 4**: Principal Component Analysis on whole genome data (bi-allelic SNPs) in A : goats, and B: sheep.

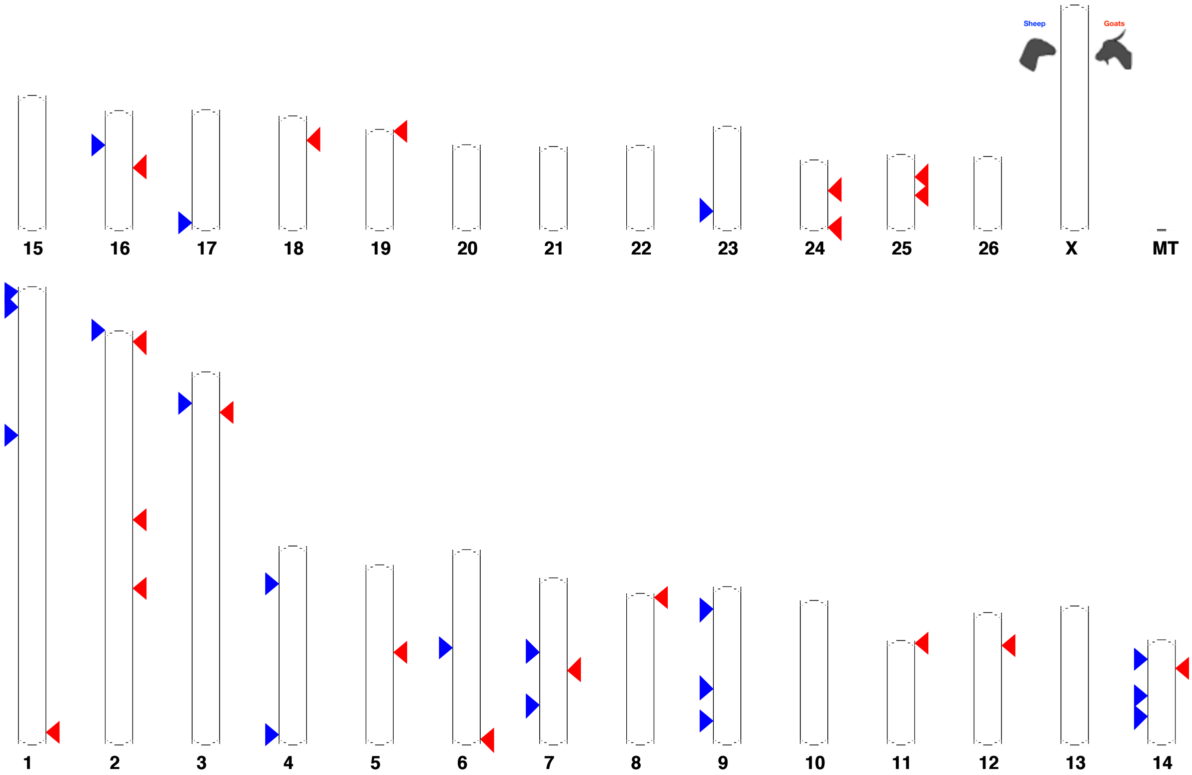


**Supp Fig 5**: Position on the sheep chromosomes of the genomic regions associated to altitude in sheep and goats .

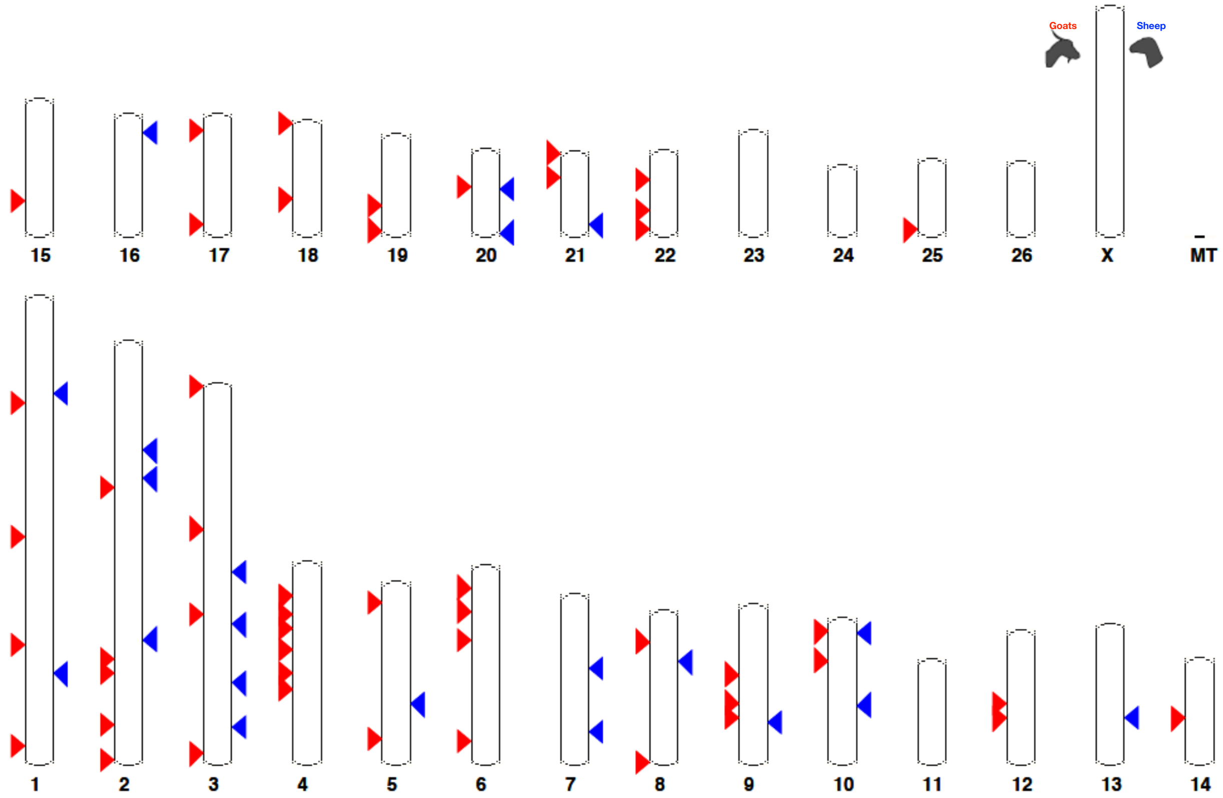


**Supp Fig 6** : Position on the sheep chromosomes of the genomic regions associated to slope in sheep and goats.

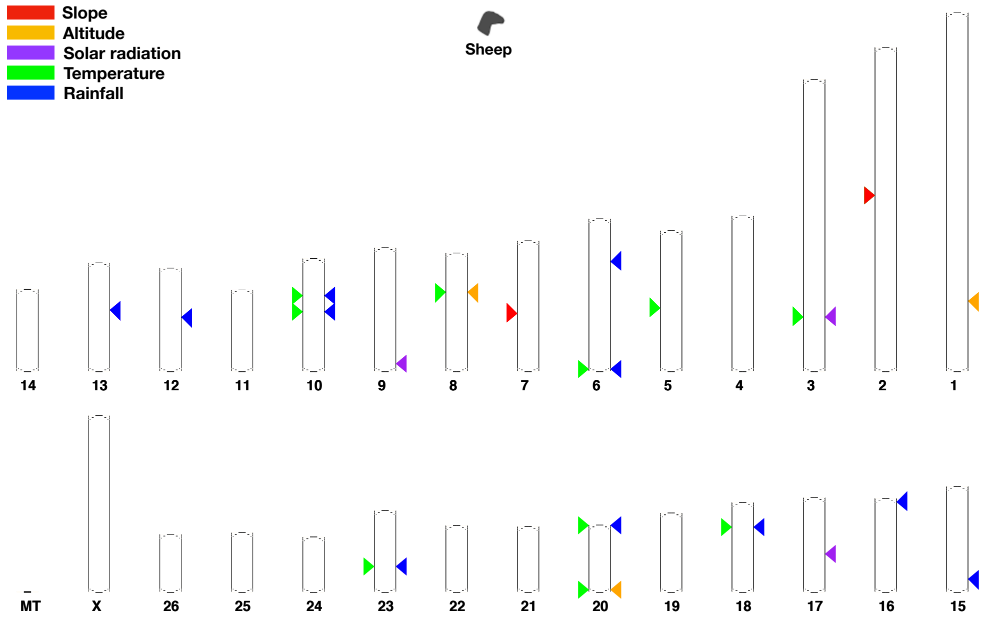


**Supp Fig 7**: Position of the major candidate genomic regions for adaptation in sheep. Displayed regions are larger than 200kb

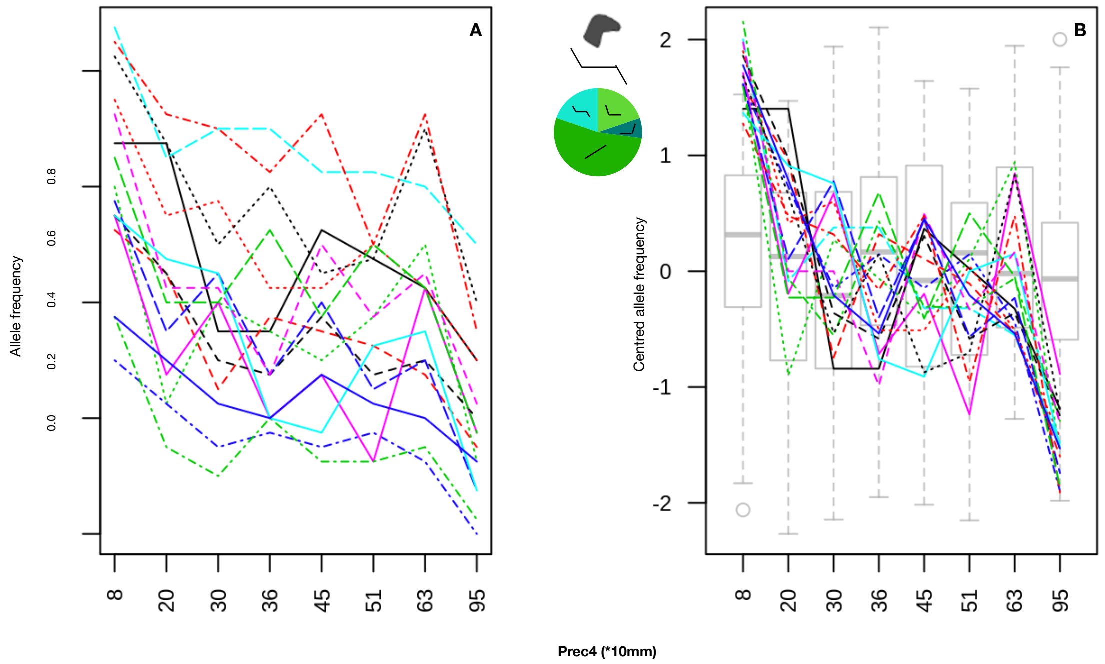


**Supp Fig 8**: Candidate variants with allele frequencies varying at both extremes of an environmental gradient (i.e., rainfall) in sheep. A: true allele frequencies. B: centred values; the grey box-plots represent neutral variations of allelic frequencies for a set of 100 random variants. The pie chart gives the proportions of variants assigned to the different types of profiles along the rainfall gradient (i.e. linear, uniform except at one extreme of the gradient, uniform except at both extremes of the gradient).

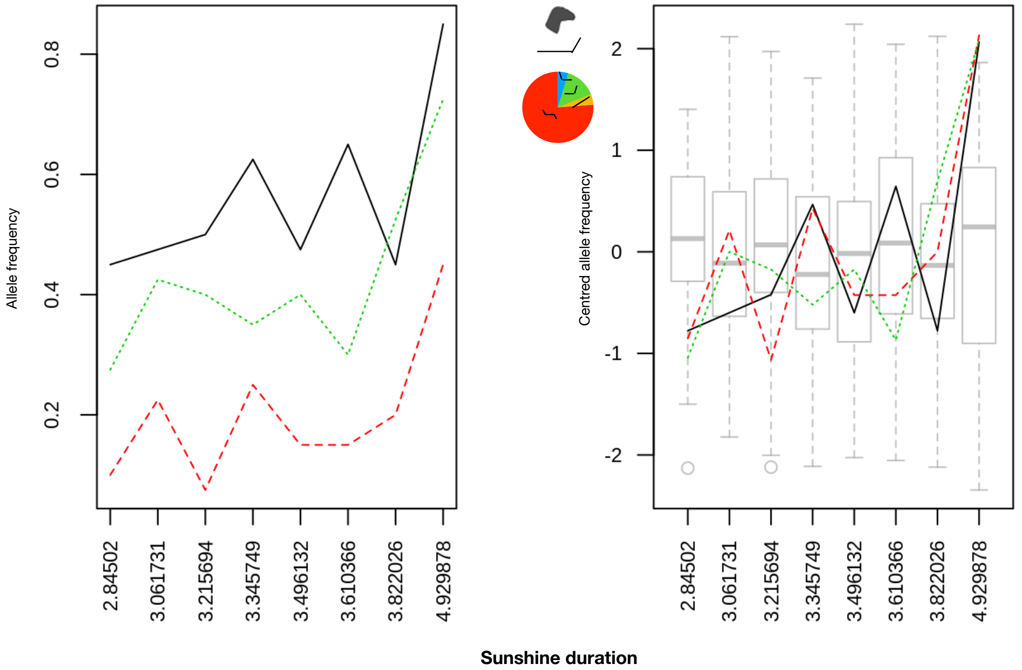


**Supp Fig 9**: Candidate variants with allele frequencies varying at one extremity of an environmental gradient (i.e., sunshine) in sheep. A: true allele frequencies. B: centred values; the grey box-plots represent neutral variations of allelic frequencies for a set of 100 random variants. The pie chart gives the proportions of variants assigned to the different types of profiles along the sunshine gradient (i.e. linear, uniform except at one extreme of the gradient, uniform except at both extremes of the gradient).

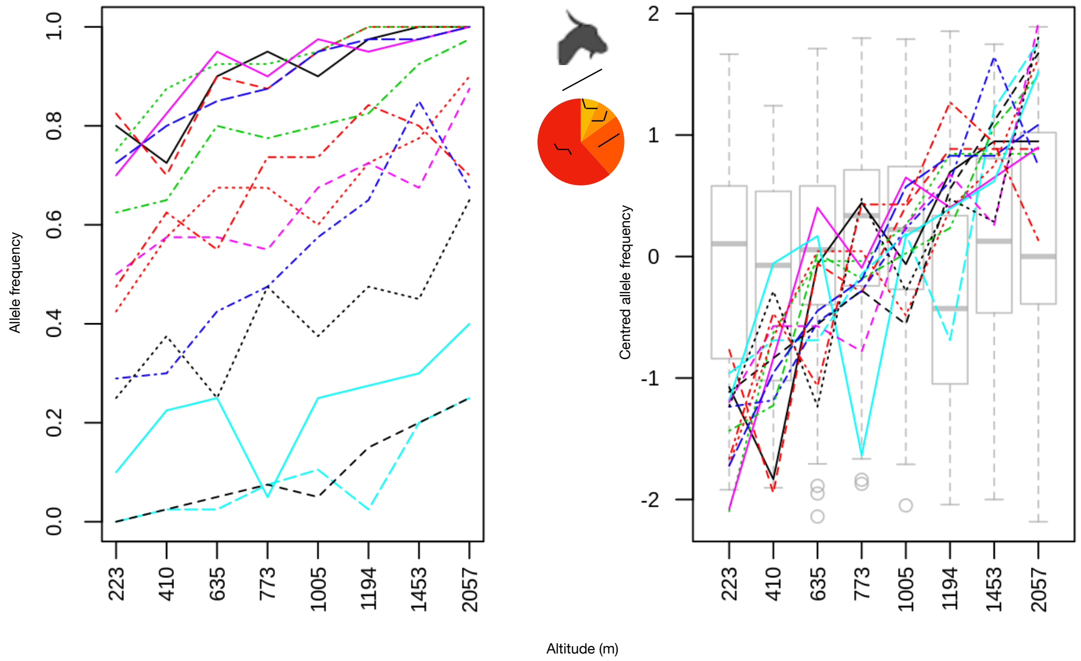


**Supp Fig 10**: Candidate variants with allele frequencies varying linearly along an environmental gradient (i.e., altitude) in goats. A: true allele frequencies. B: centred values; the grey box-plots represent neutral variations of allelic frequencies for a set of 100 random variants. The pie chart gives the proportions of variants assigned to the different types of profiles along the altitude gradient (i.e. linear, uniform except at one extreme of the gradient, uniform except at both extremes of the gradient).

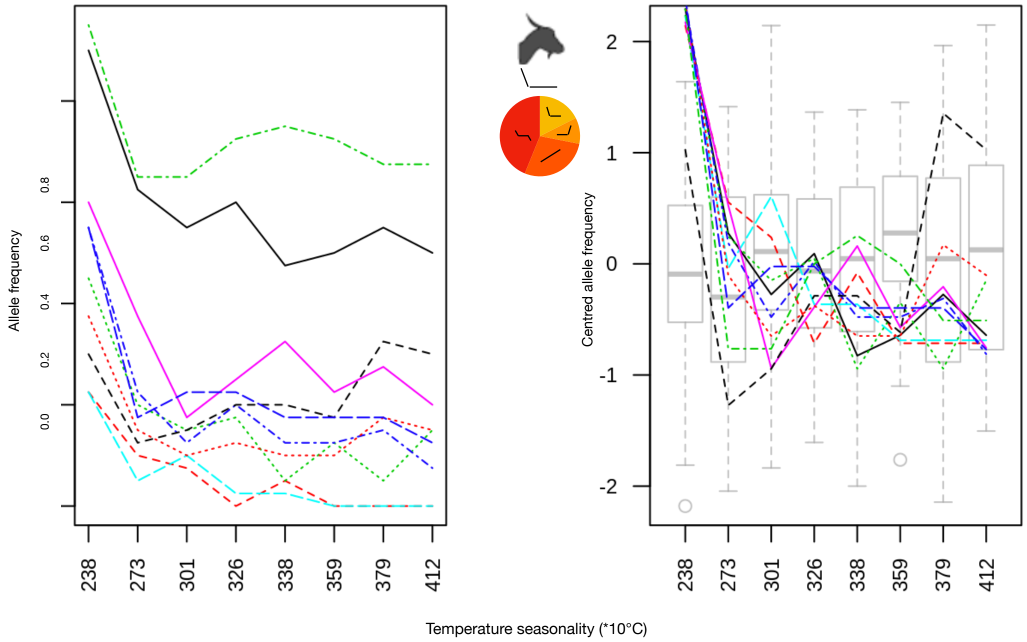


**Supp Fig 11**: Candidate variants with allele frequencies varying at one extremity of an environmental gradient (i.e., temperature seasonality) in goats. A: true allele frequencies. B: centred values; the grey box-plots represent neutral variations of allelic frequencies for a set of 100 random variants. The pie chart gives the proportions of variants assigned to the different types of profiles along the temperature seasonality gradient (i.e. linear, uniform except at one extreme of the gradient, uniform except at both extremes of the gradient).

**Supp Fig 12**: Manhattan plots of Q-values obtained from Samβada's G scores. Only plots with significant associations are shown for both sheep and goats. False discovery rate, represented by the horizontal red line, was set at 10%.

Sheep tmean7

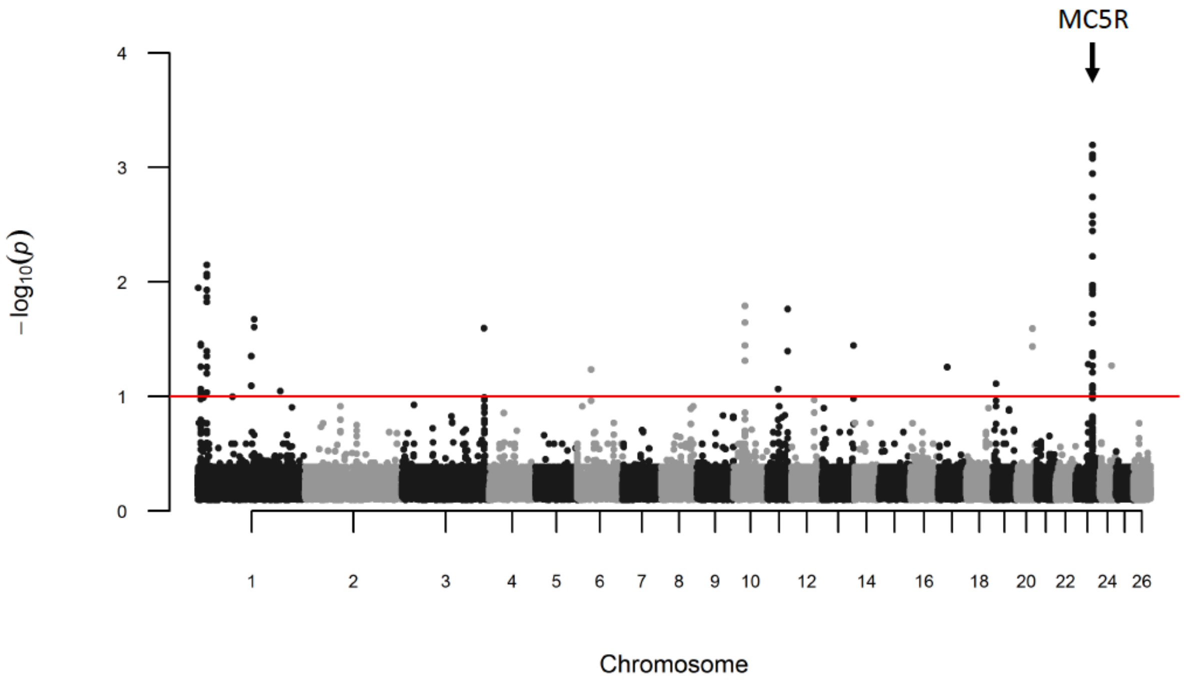


Sheep prec4

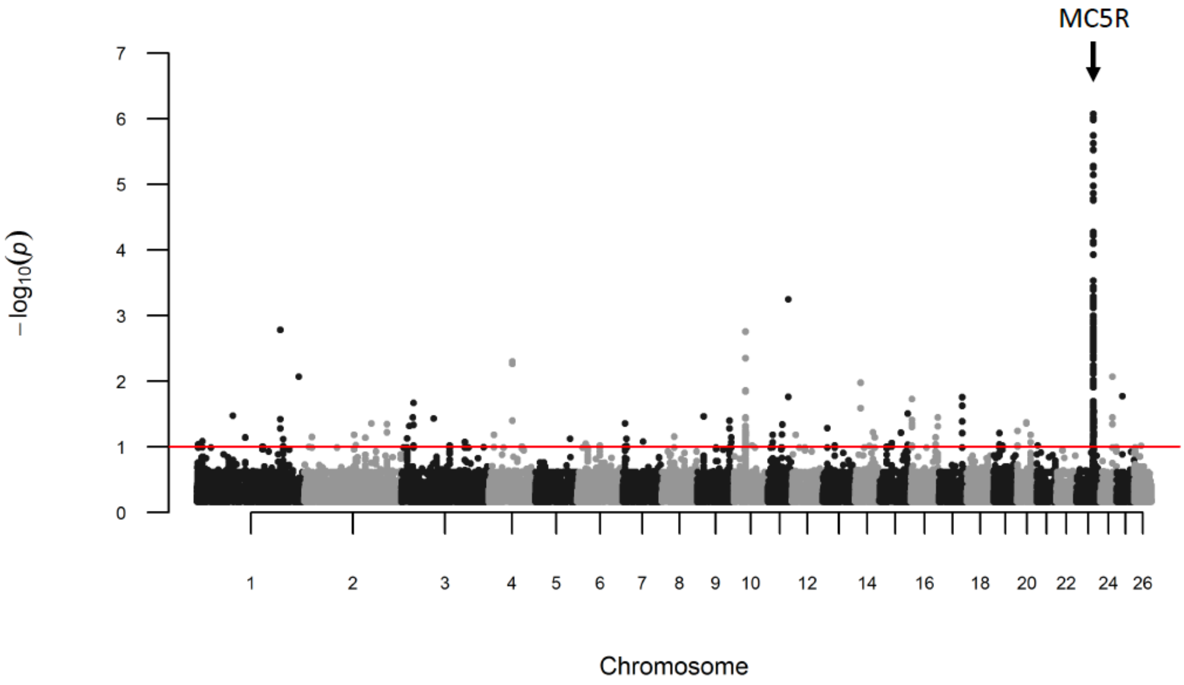

**Supp Fig 12 continued**

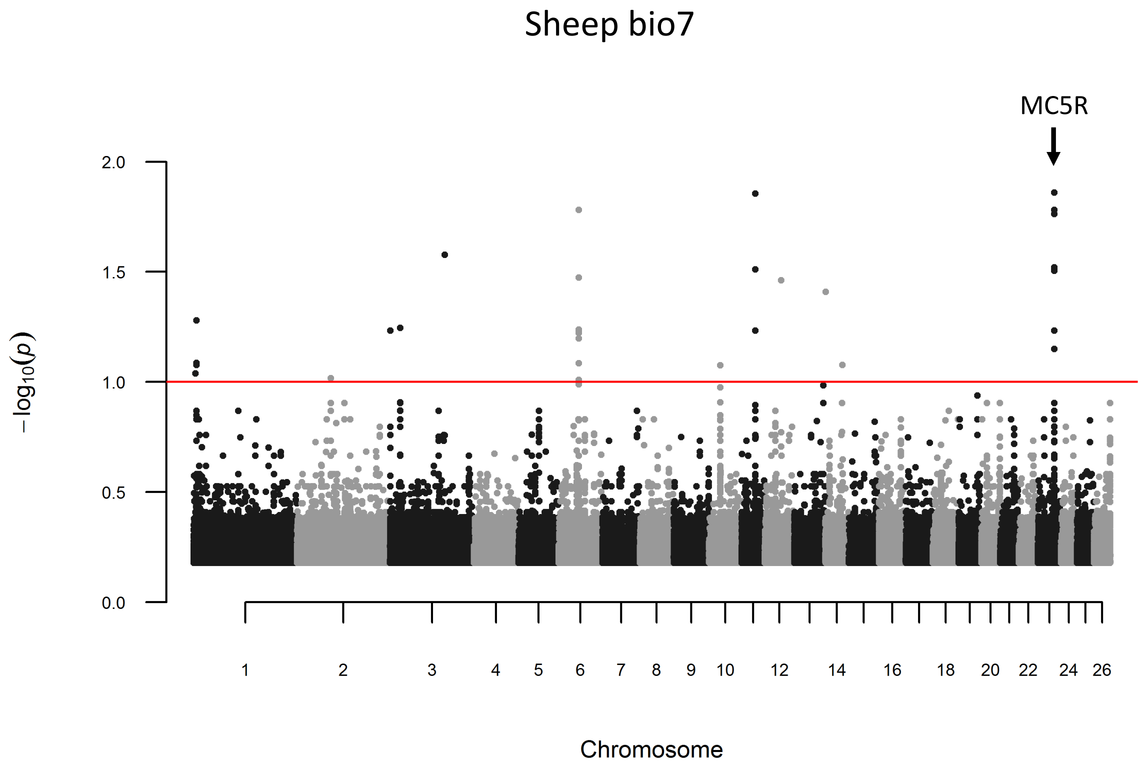


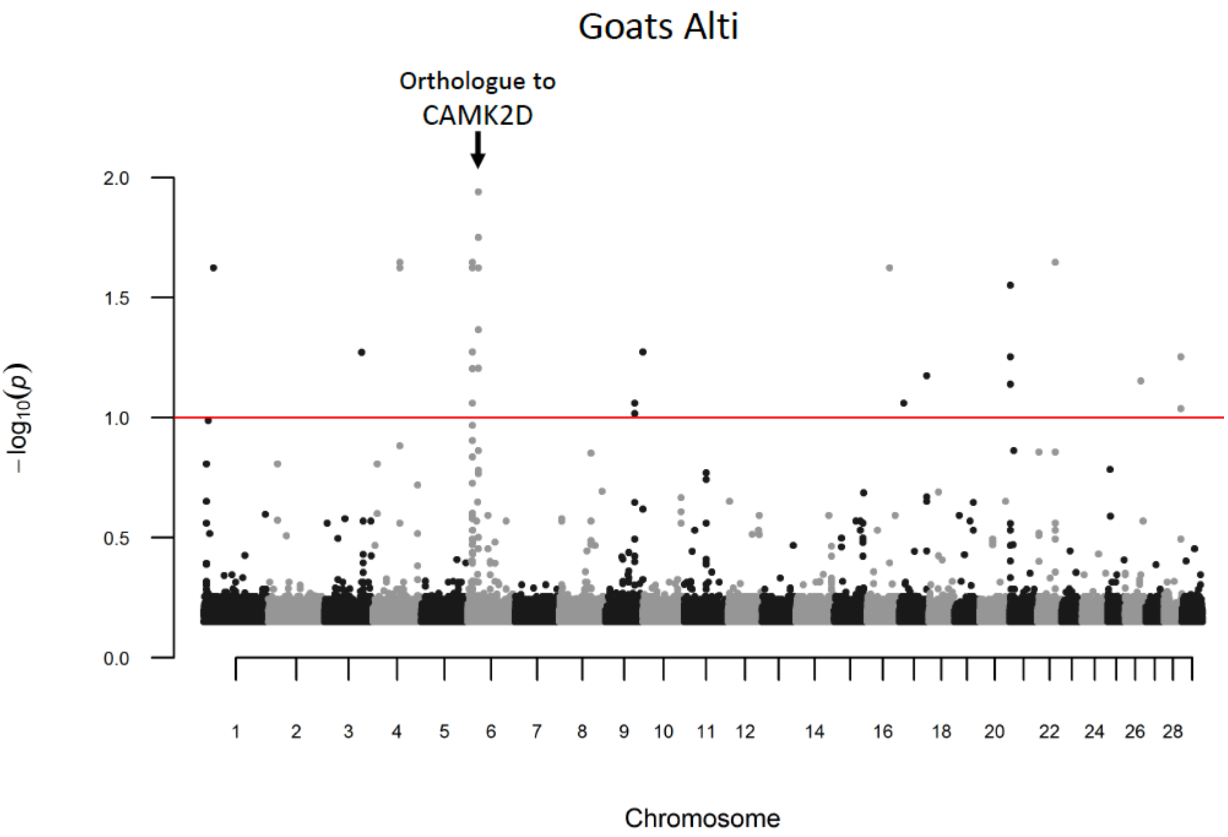

**Supp Fig 12 continued**

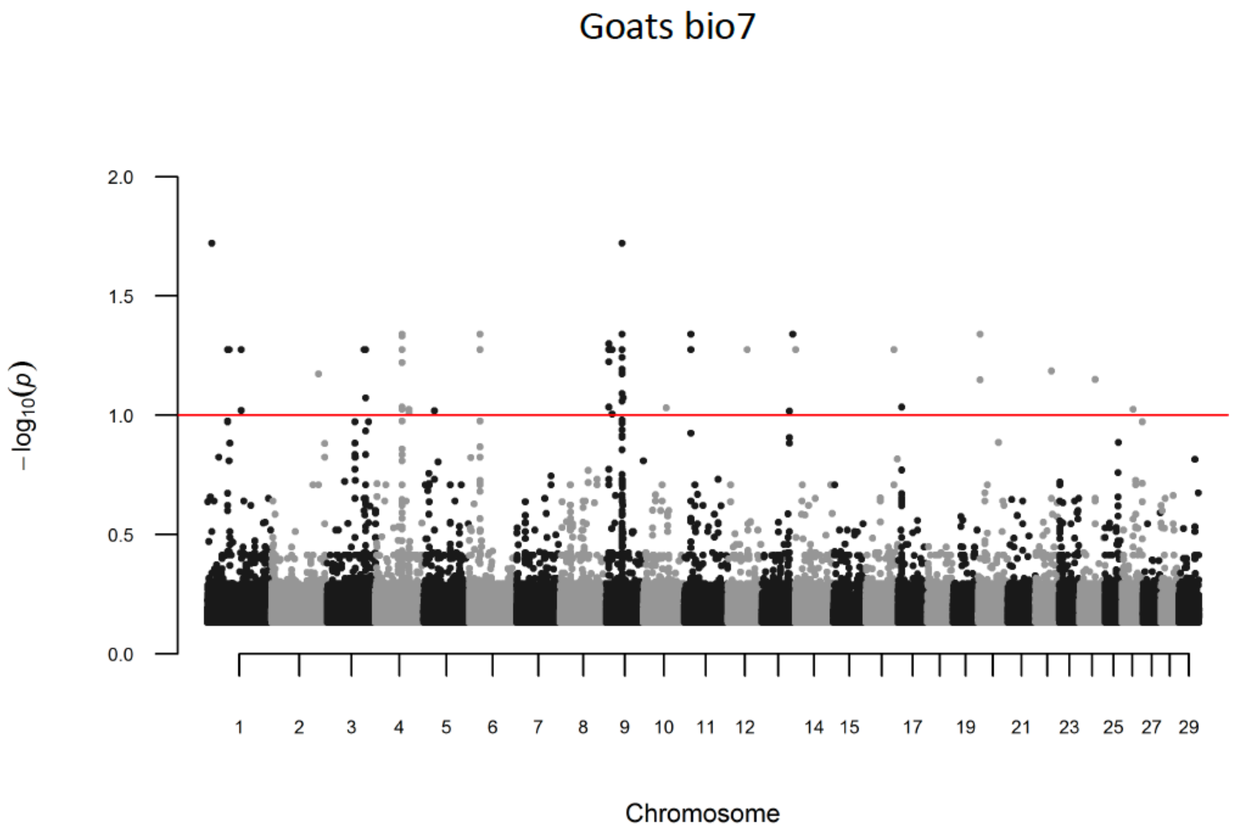

### Goats bio15

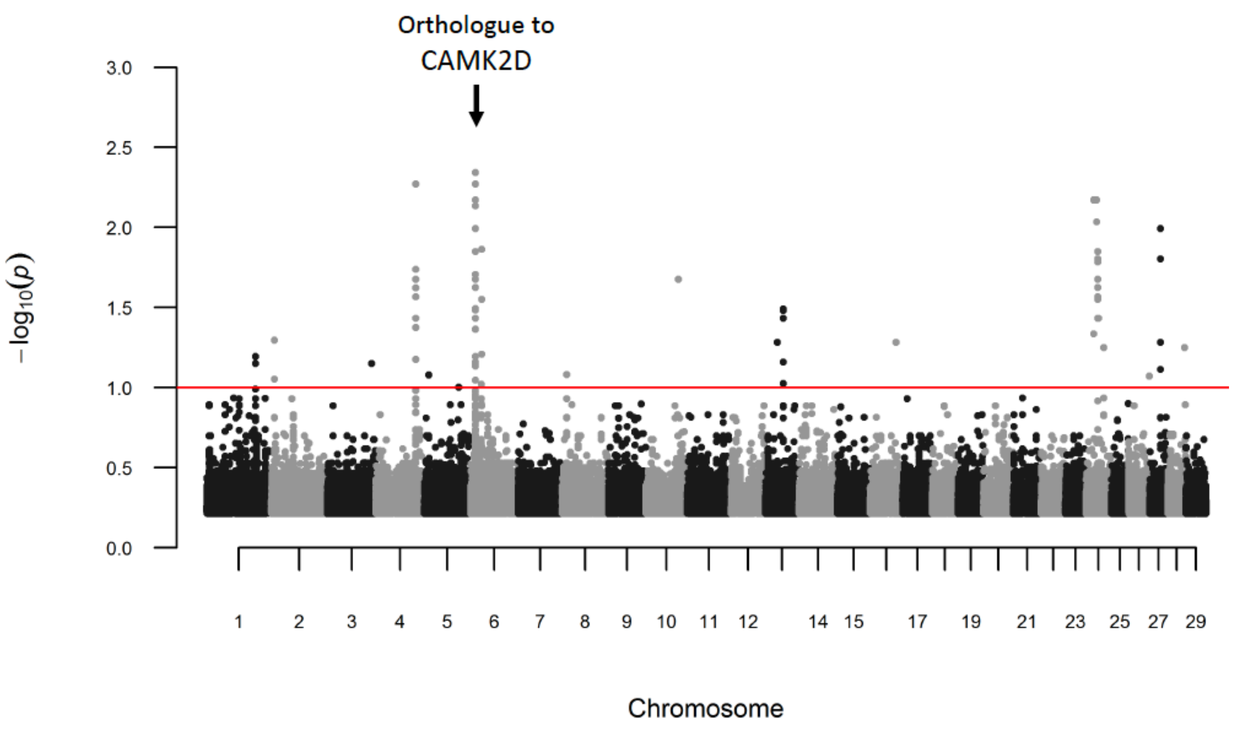

Orthologue to
CAMK2D
3.0
2.5
2.0
1.5
1.0
0.5
0.0
$-\log_{10}(p)$
1
2
3
4
5
6
7
8
9
10
11
12
14
15
17
19
21
23
25
27
29
Chromosome

**Supp Fig 13:** Plot of XP-CLR scores along autosomes in selective sweep analysis. The horizontal red lines indicate 0.01% autosomal-wide cut-off level. The species and the environmental variable considered are displayed in the top of each graphic.

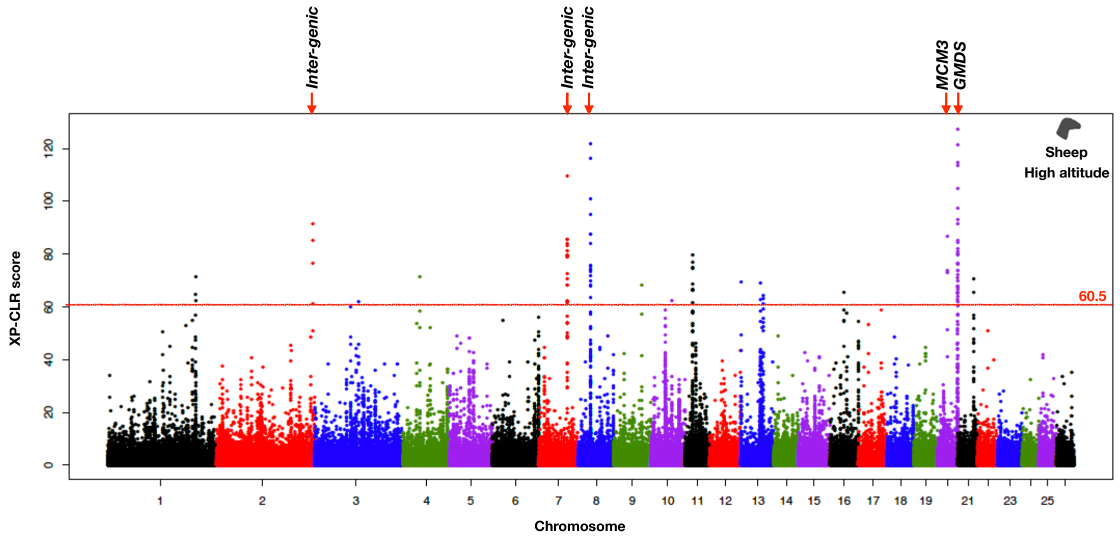


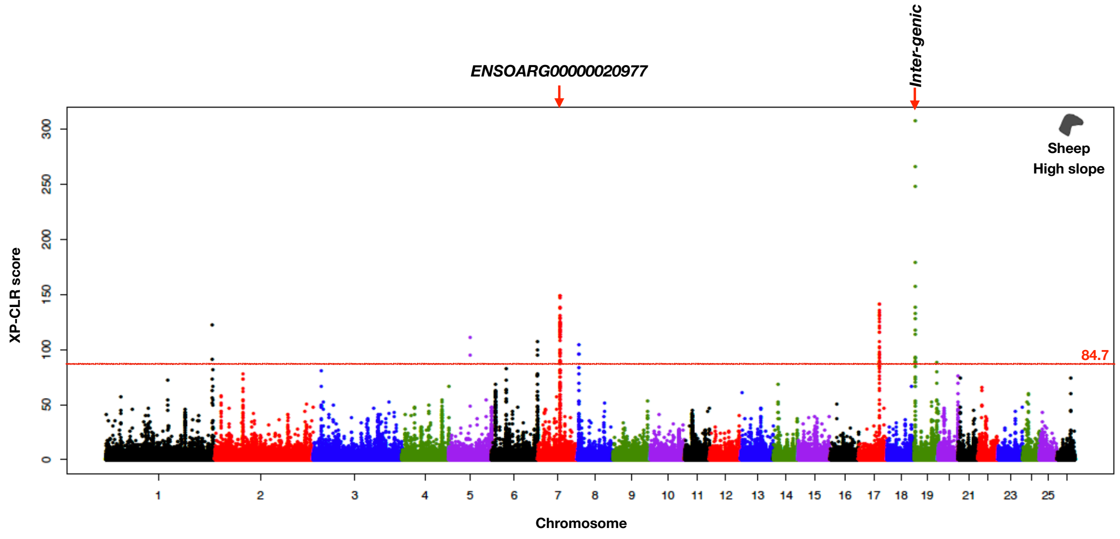

### Supp Fig 13 Continued

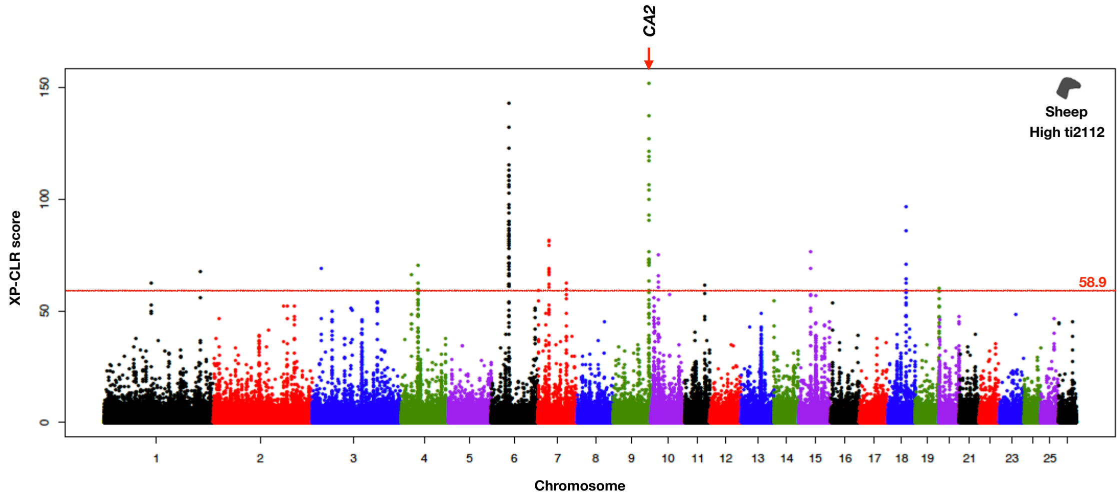


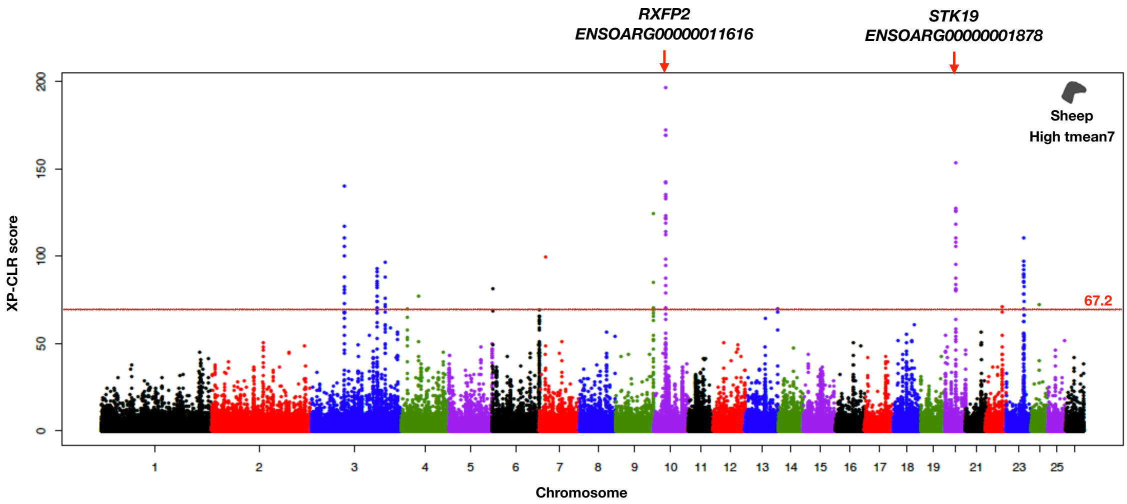

#### Supp Fig 13 Continued

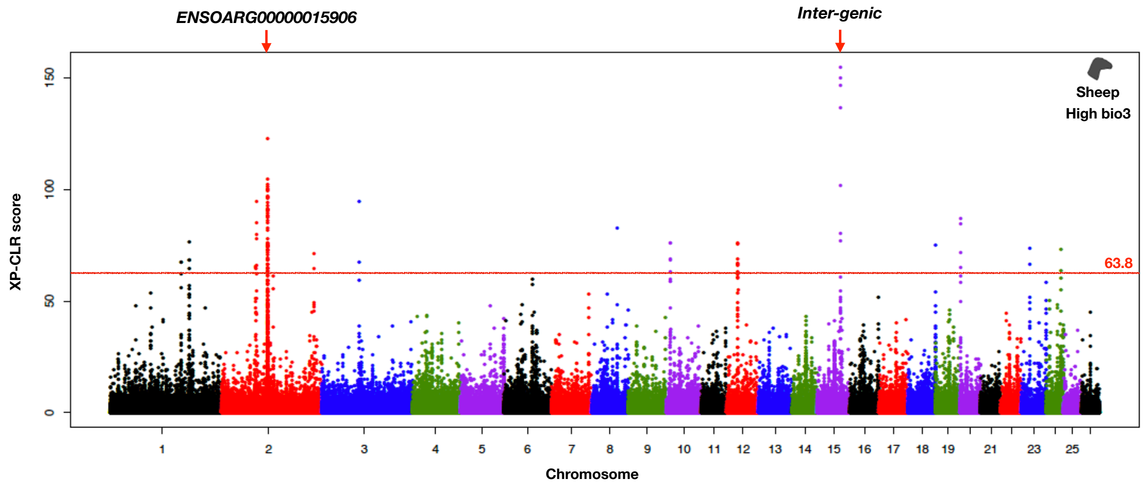


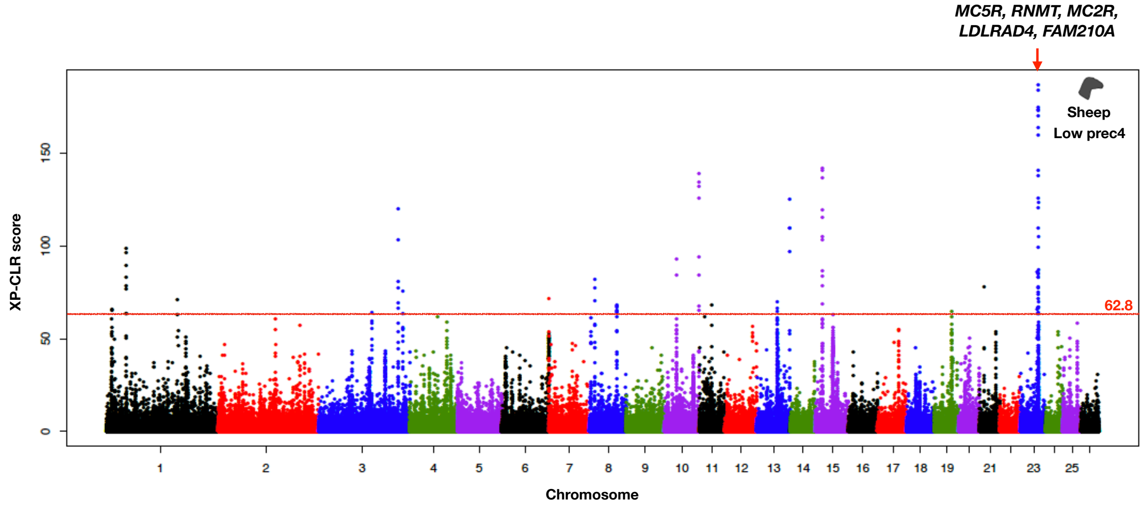

#### Supp Fig 13 Continued

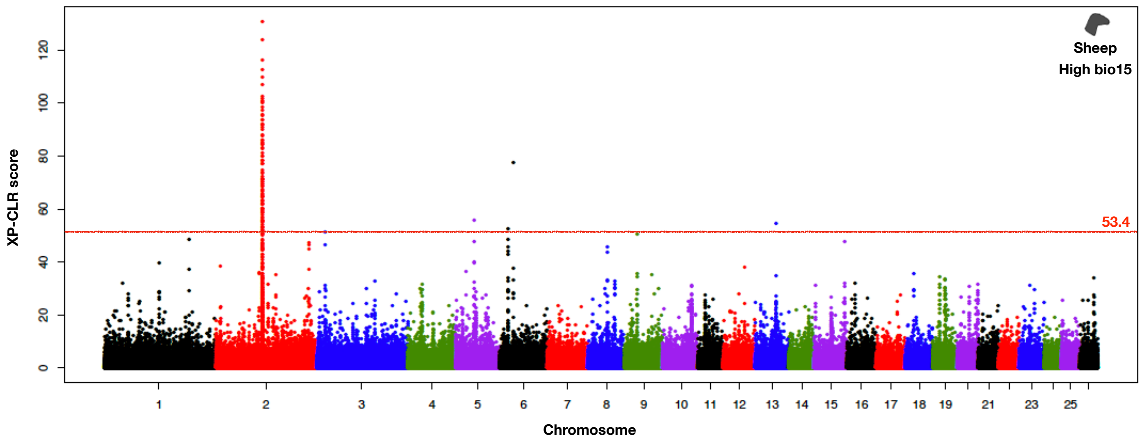


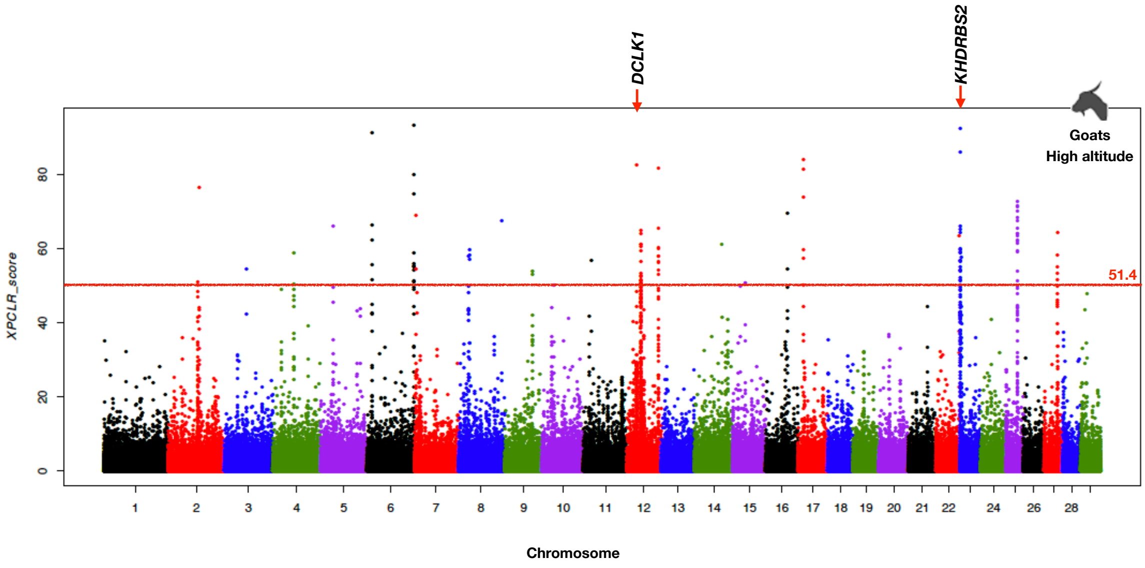

### Supp Fig 13 Continued

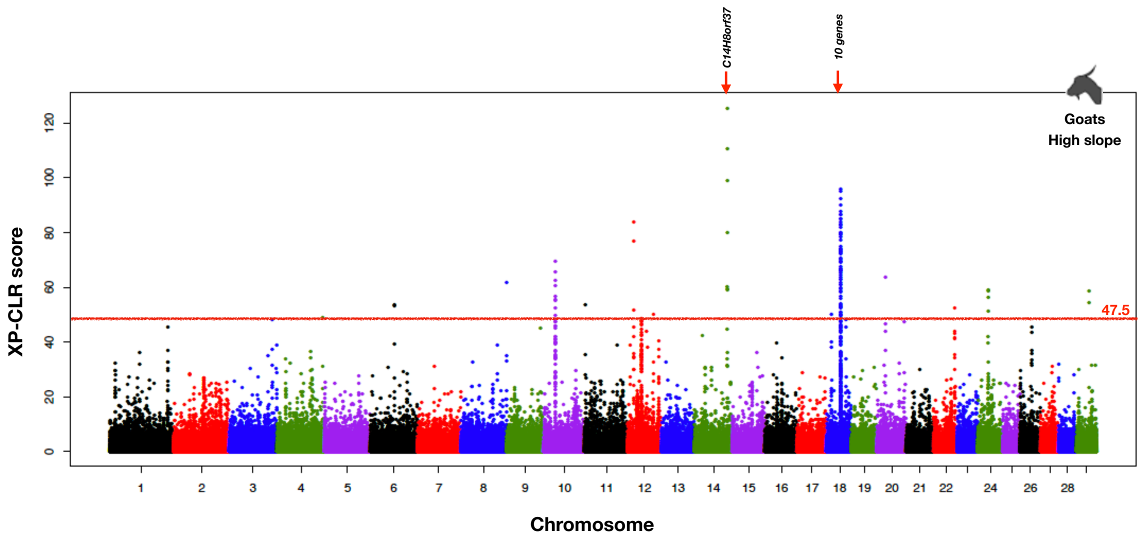


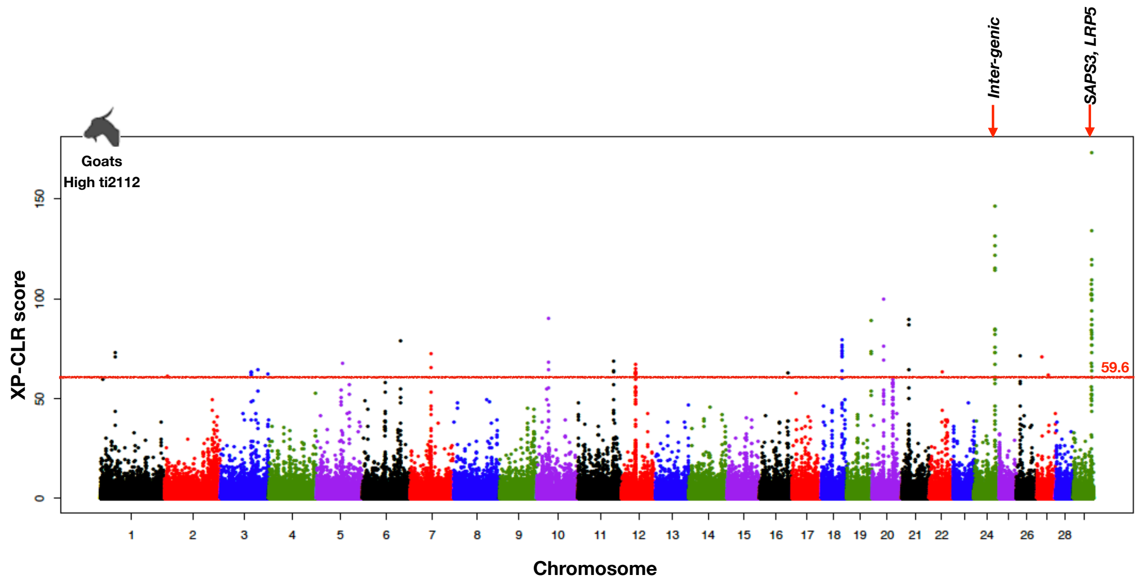

#### Supp Fig 13 Continued

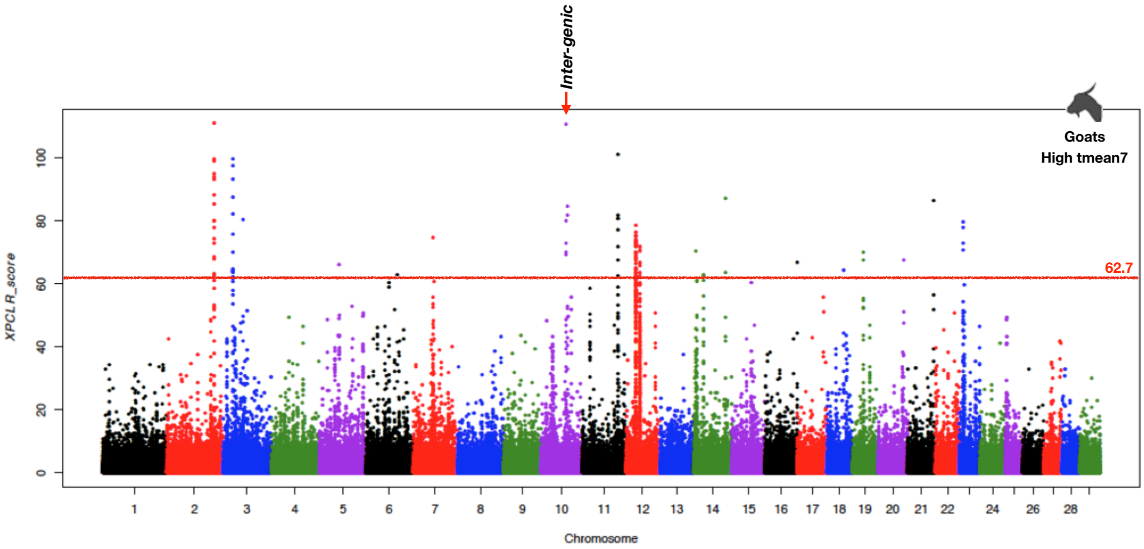


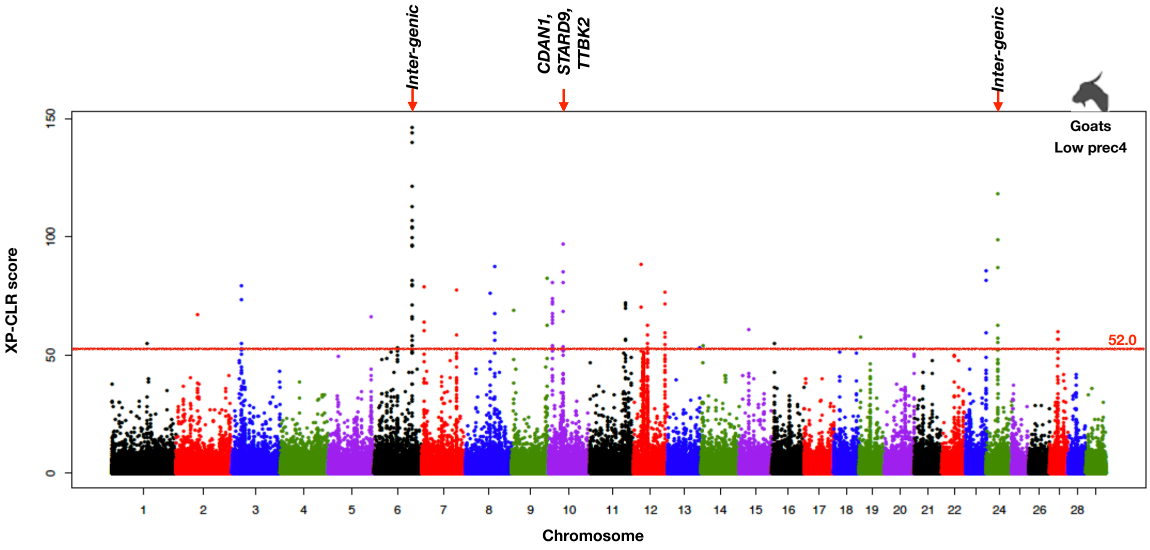

#### Supp Fig 13 Continued

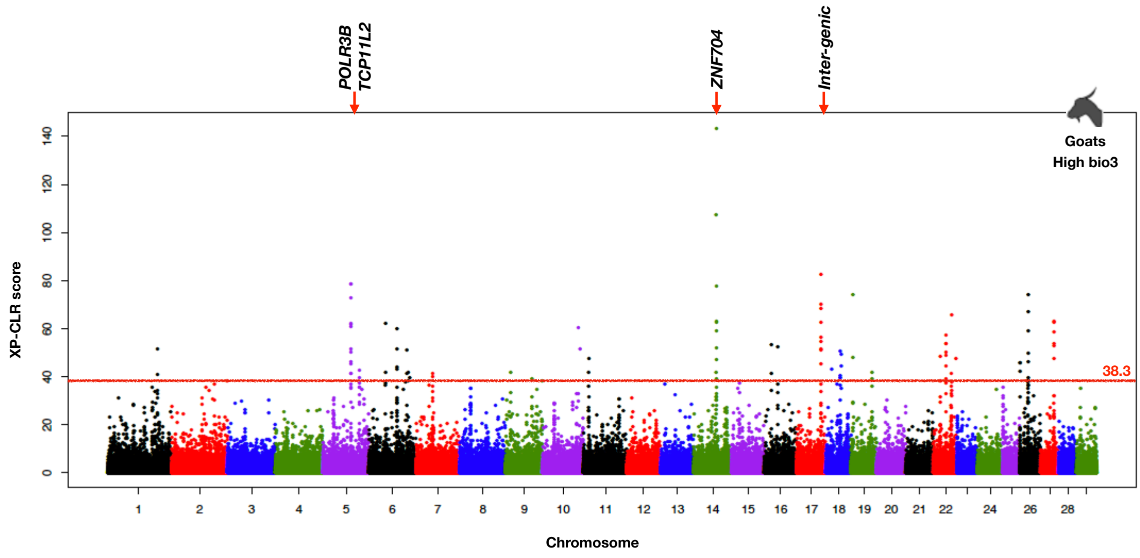


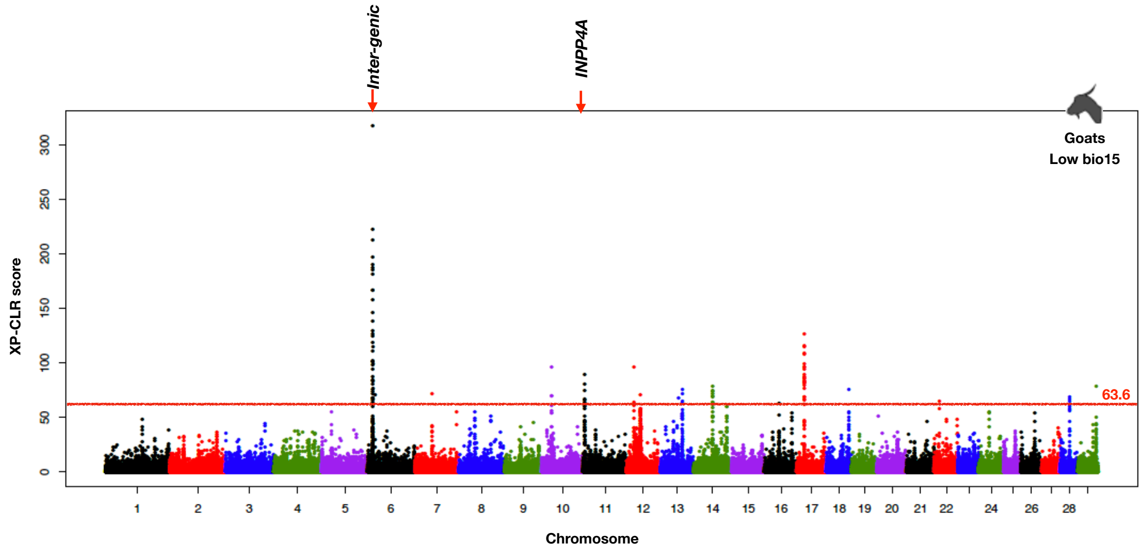
